# Nesprin-2 accumulates at the front of the nucleus during confined cell migration

**DOI:** 10.1101/713982

**Authors:** Patricia M. Davidson, Aude Battistella, Théophile Déjardin, Timo Betz, Julie Plastino, Nicolas Borghi, Bruno Cadot, Cécile Sykes

## Abstract

The mechanisms by which cells exert forces on their nuclei to migrate through openings smaller than the nuclear diameter remain unclear. In microfluidic devices, the hourglass shape of the nucleus and its strain patterns as it translocates through narrow constrictions suggest pulling forces. We use CRISPR/Cas9 to fluorescently label nesprin-2 giant, a protein that links the cytoskeleton to the interior of the nucleus. We demonstrate that nesprin-2 giant accumulates at the front of the nucleus during nuclear deformation through narrow constrictions, independently of the nuclear lamina. We find that nesprins are more mobile than lamin A/C, at time scales similar to that of the accumulation. Using artificial constructs, we show that the actin-binding domain of nesprin-2 is necessary and sufficient to generate this accumulation, and that microtubules are not necessary. Actin filaments are organized in a barrel structure around the moving nucleus in the direction of movement, suggesting that this structure is responsible for redistribution of nesprins towards the front of the nucleus. Two-photon ablation and the use of drugs inhibiting the cytoskeleton demonstrate a pulling force on the nucleus from the front of the cell that is dependent on formin and actomyosin contractility. This elastic recoil is significantly reduced when nesprins are reduced at the nuclear envelope. We thus show that actin redistributes nesprin-2 giant towards the front of the nucleus and contributes to pulling the nucleus through narrow constrictions, in concert with myosin.

## INTRODUCTION

Mammalian cells migrate through tissues during essential processes, including wide-scale migration throughout development,^1^ immune cell migration to sites of inflammation and fibroblast migration to repair wounds^2^. Migration also occurs in disease, for example during cancer cell invasion to establish distant metastatic tumors.^3^ These cells must crawl through narrow openings in cell-cell junctions, the extracellular matrix and basement membranes. In most cells, nuclear deformability is the limiting factor for migration through constrictions smaller than the diameter of the nucleus.^4,5^ The mechanisms by which cells can exert forces to deform their nuclei remain unclear.^6^

A mechanical link between the nucleus and the cytoskeleton is provided by nesprins. They are a family of nuclear envelope proteins (nesprins1-4 and KASH5) defined by their Klarsicht/ANC-1/Syne homology (KASH) domain.^7^ This domain binds to SUN proteins in the perinuclear luminal space; SUN proteins cross the inner nuclear membrane and bind to lamins at the nuclear interior. By crossing the outer nuclear membrane, nesprins provide a mechanical link from the cytoplasm to the nuclear interior. The largest isoforms of nesprins-1 and 2 are termed “giant” due to their size (approximately 1 MDa and 800 kDa)^8^. They consist of a KASH domain and an N-terminal calponin homology (actin-binding) domain separated by 74 and 56 spectrin repeats, respectively. The genes that encode these giant nesprins (SYNE1 and SYNE2) can also give rise to shorter isoforms that may or may not contain the KASH or actin-binding domains.^8^ Nesprin-2 can also indirectly bind actin through fascin^9^ and FHOD1^10^, possibly reinforcing its mechanical link to actin. Nesprin-1 and nesprin-2 bind microtubules through kinesin-1 and dynein^11^ and plectin can link another nesprin, nesprin-3, to the intermediate filament vimentin.^12^

Nesprins are implicated in nuclear movement and positioning in fibroblasts and muscle cells^13^, outer hair cells^14^, and during retinal and neuronal cell development.^11,15^ In particular, nesprin-2 giant determines nuclear re-centering after displacement due to centripetal forces in NIH 3T3 fibroblasts (that do not express nesprin-1 giant)^16^. Nesprin-2 also enables migration through 5 μm constriction and restrictive collagen gels, in concert with non-muscle myosin IIb, an actin motor protein.^17^ Cells lacking fascin, which provides additional links between actin and nesprin-2, have defects in translocating their nuclei through narrow constrictions.^9,18^

Accumulation of non-muscle myosin IIb at the rear of the cell as the nucleus is squeezed through a constriction indicates actomyosin contractility pushes the nucleus forward.^6^ However, the hourglass shape of fibroblast nuclei deforming through microfluidic constrictions and the patterns of strain in the nuclear interior suggests a pulling force from the front of the nucleus as the origin of nuclear motion.^19^ In cells undergoing lobopodial migration, inhibition of actomyosin contractility at the front of the cell results in the nucleus falling back, again pointing to a forward pulling mechanism.^20^

Previous studies of giant nesprins relied on protein knock-down or mini-nesprins constructs.^21^ These artificial constructs were conceived to recapitulate the link between the nucleus and actin provided by nesprin-2 in a short construct that is easily expressed. They consist of the actin-binding and nucleus-anchoring domains of nesprin-2 giant, along with only 4 of the 56 spectrin repeats. These studies have evidenced the links to SUN proteins^21^ and transmembrane actin-associated nuclear (TAN) lines^22^, which couple nesprin-2 to actin filaments at the dorsal side of the nucleus. However, the spectrin repeats that are left out may provide important sites of interaction. Notably, domains that bind to cytoskeletal filaments, motor proteins and proteins that cross-link the cytoskeleton to nesprins are not present in these artificial constructs.^9,10,12^ To obtain a more complete picture of the mechanical link between the giant nesprin and the cytoskeleton we employ CRISPR to tag the endogenous protein.

We show that the actin-binding domain of nesprin-2 is necessary and sufficient to generate accumulation of nesprin-2 giant at the front of the nucleus during migration through narrow microfluidic constrictions. This mechanism is independent of A-type lamins and microtubules. We demonstrate a barrel shape of filamentous actin around the nucleus, which colocalizes with nesprin-2, but strikingly actin is not enriched at the front surface of the nucleus. Our results suggest that this barrel structure has a role in nesprin distribution. In ablation experiments we find that the nucleus is under pulling forces that are reduced when nesprins are displaced from the nuclear envelope. Altogether our results indicate that actin is involved in redistributing nesprin-2 towards the front of the nucleus and that the nucleus is pulled through constrictions via nesprins.

## RESULTS

### Cell migration through constrictions causes nesprin-2 accumulation at the front of nuclei

Inspired by previous work using mini-nesprin constructs,^21^ we added a green fluorescent protein (GFP) sequence to the N-terminus of endogenous *Syne2* by CRISPR-Cas9, immediately preceding the sequence that encodes the actin-binding domain of nesprin-2 (Figure 1A and Supplemental Figure 1A). Here we use mouse embryonic fibroblasts (MEF), that express nesprin-2 giant but not nesprin-1 giant,^22^ as validated by qPCR (Supplemental Figure 1B). After validation by PCR, sequencing and Western Blotting (Supplemental Figure 1C, 1D) we selected a clonal cell population with a homozygous modification. By epifluorescence microscopy we detected a green signal around the periphery of the nucleus (Figure 1B), consistent with the known localization of nesprin-2 giant at the outer nuclear membrane. We hereafter call the modified protein “GFP-nesprin-2”.

**Figure 1.**
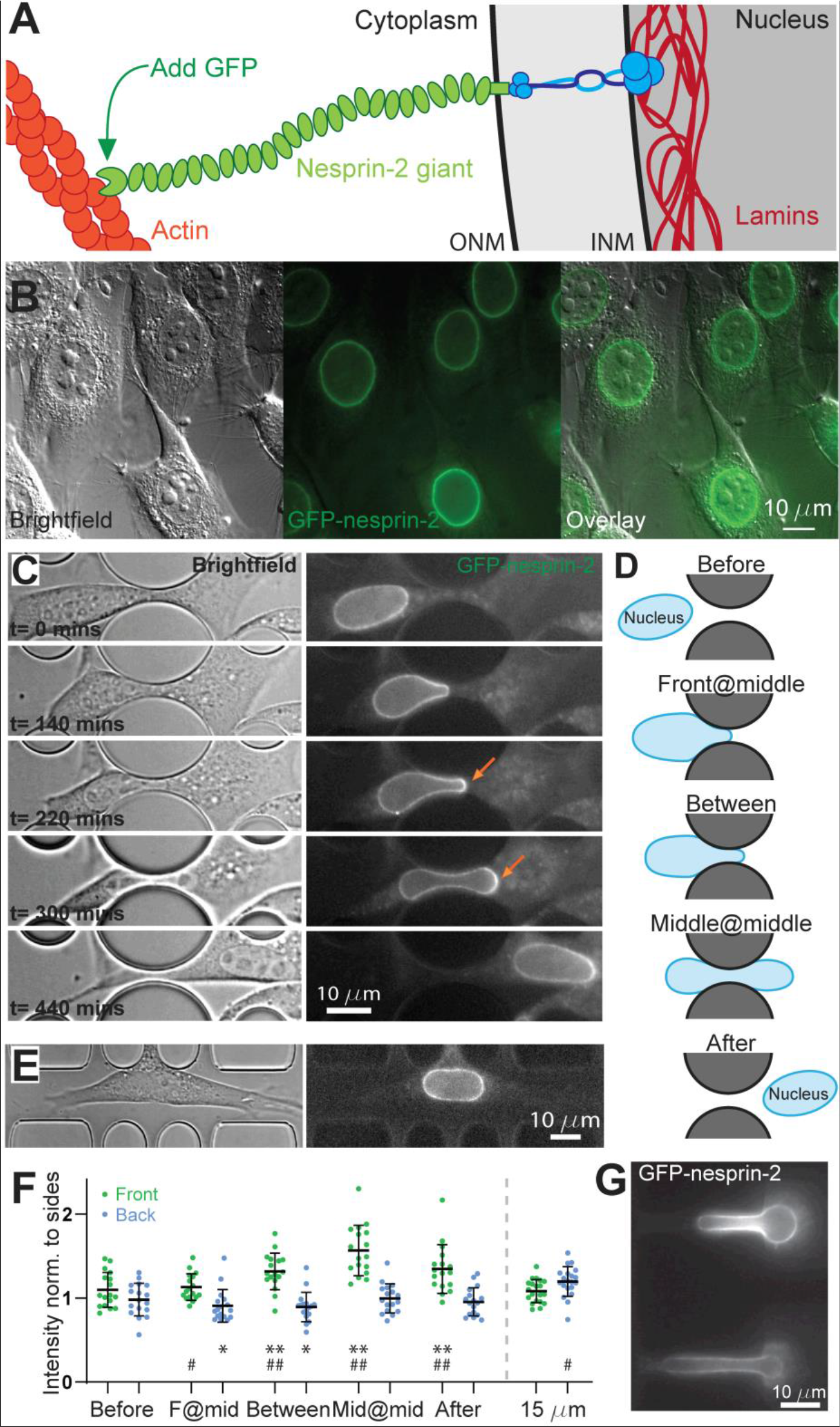
Giant nesprins accumulate at the front of the nucleus during migration through narrow constrictions. **A**, Schematic illustration of the nuclear envelope, depicting the link between the lamina, SUN proteins, giant nesprins and actin. **B**, Mouse embryonic fibroblasts genetically modified using CRISPR to introduce a GFP sequence immediately after the start codon of *Syne2* display fluorescence at the nuclear envelope. **C**, Brightfield and fluorescent images of a cell migrating through a 2 μm constriction. Arrows indicate areas of accumulation of nesprin-2 at the front of the nucleus as it is deformed through the constriction. (See also movie 1.) **D**, Schematic illustrating the 5 time points studied in the analysis: Before nucleus deformation (“Before”), once the front of the nucleus reaches the middle of the constriction (“Front@mid”), once the middle of the nucleus reaches the middle of the constriction (“Middle@middle”), a timepoint halfway between Front@mid and Middle@middle (“between”), and late time point after the deformation (“After”). **E**, Brightfield and fluorescent images of a cell migrating along a 15 μm control channel. **F**, Quantification of the intensity of the fluorescent signal around the perimeter of the nucleus in a 2 μm constriction and a 15 μm control channel. The average intensity at the front and the back is normalized to the intensity at the sides, demonstrating an increase in the signal at the front of the nucleus as it is deformed and a reduction after deformation. (Error bars correspond to the standard deviation. Constriction: n=17 cells over 4 different experiments, 15 μm channels: n=20 cells over 3 experiments. p-values: *: p<0.05 compared to Before, **: p<0.01 compared to Before, # p<0.05 compared to side, ##: p<0.01 compared to side) **G**, GFP-nesprin-2 cells deformed by the pressure gradient in a micropipette aspiration device. Note that the GFP signal does not increase at the tip but increases at sides of the nucleus.

We characterize the localization of nesprin-2 during cell migration through narrow constrictions using migration devices^4,23^ consisting of pillars delimiting control channels and constrictions smaller than the nucleus diameter (Supplemental Figure 1E). When MEFs migrate through 2 μm constrictions they extensively deform their nuclei (Figure 1C)^4^ and the GFP signal increases at the front of the nucleus (orange arrows in Figure 1C). To quantify this accumulation, we measure the intensity around the periphery (Supplemental Figure 1F) of several nuclei at five time points during deformation (Figure 1C,D,F) and in non-constricting channels (Figure 1E,F). Indeed, the intensity at the front of the nucleus increases relative to the sides as the nucleus deforms through the constriction, peaking as the middle of the nucleus reaches the middle of the constriction (Figure 1C and F). We confirmed this accumulation in immunofluorescence measurements (Supplemental Figure 1G). Intensity at the front increases with time at a slower rate than it decreases once the nucleus has passed through the constriction (Supplemental Figure 1H). Notably, in the 15 μm control channel, there is an increase in intensity at the front although this effect is lower than in constrictions.

To determine whether the accumulation of nesprin-2 at the front could be due to a change in nucleus shape, we deform GFP-nesprin-2 cells in micropipette aspiration devices (Figure 1G).^24^ Cells deformed under such exogenous forces display a completely different distribution of nesprins: we observe a decrease in signal at the tip of the nucleus and an increase at the sides (Figure 1G). This accumulation is likely due to stretching of the membrane at the tip or accumulation of the membrane at the edges (see below). Nesprin accumulation at the front of the nucleus thus results from active migration through narrow constrictions rather than passive nucleus deformation by exogenous forces.

### Nesprin-2 accumulation does not associate with lamin A/C accumulation

As nesprin-2 accumulation results from an active process of migration through a narrow constriction, we sought to determine whether it could cause or be caused by nuclear lamina remodeling. To do so, we fluorescently tagged A-type lamins in GFP-nesprin-2 cells. To avoid lamin overexpression (which could reduce nuclear deformability)^25^ we created a cell line, using CRISPR-Cas9, that expresses a red fluorescent protein (mCherry) coupled to endogenous lamin A/C. We call this modified protein “mCh-LAC”. After clonal selection and validation, we obtained a heterozygous cell line (Supplemental Figure 2A and B).

**Figure 2.**
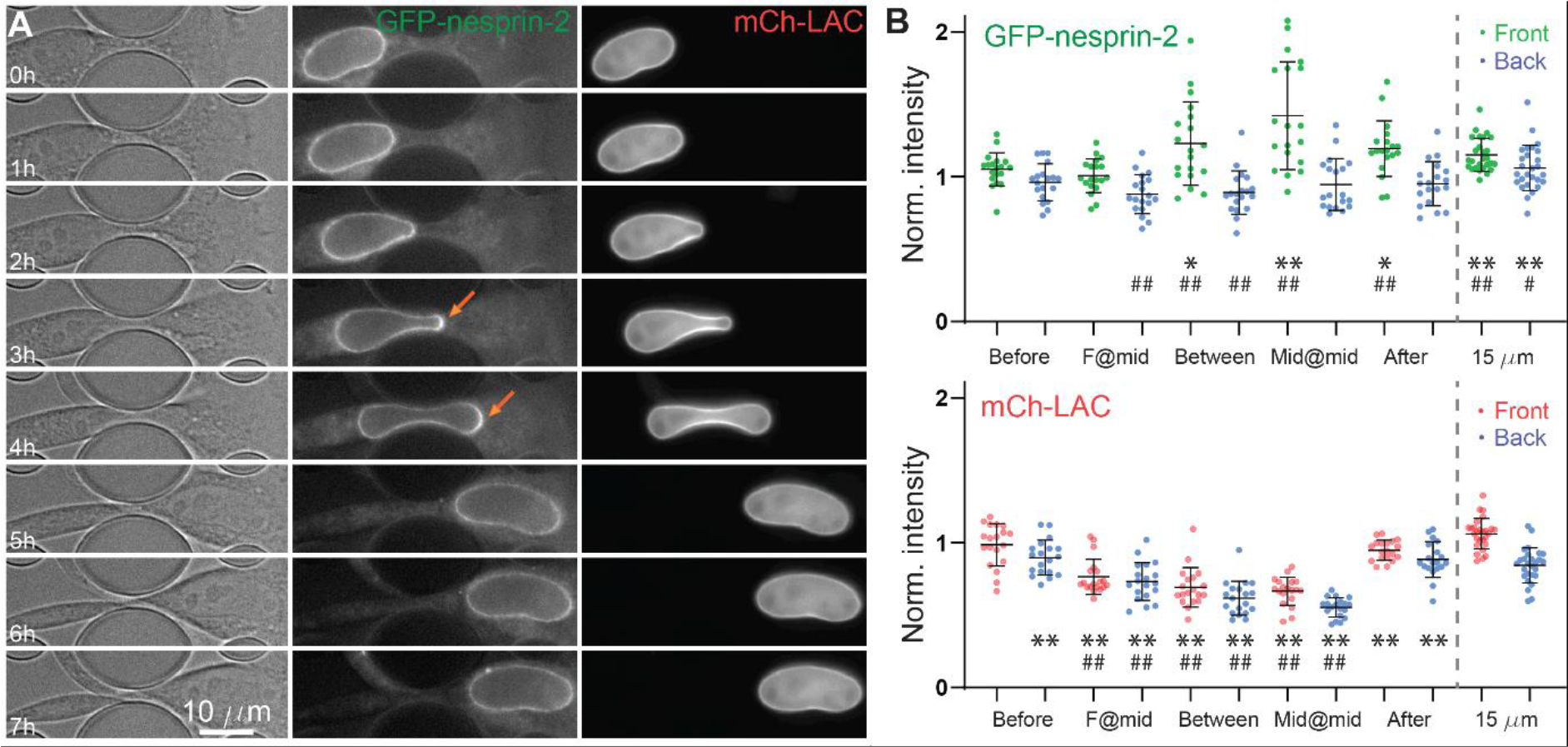
The accumulation of nesprins does not associate with lamin accumulation. **A**, CRISPR-modified cells (Brightfield, left) that express GFP-nesprin-2 (middle) and mCh-LAC (right) show nesprin but not lamin accumulation at the front of the nucleus as it is deformed through a narrow constriction. (See also movie 2.) **B**, Quantification of intensity of GFP-nesprin-2 and mCh-LAC, normalized to the intensity at the sides. (Error bars correspond to the standard deviation. Constriction: n=19 cells over 5 different experiments, 15 μm channel: n=27 over 6 experiments. p-values: *: p<0.05 compared to Before, **: p<0.01 compared to Before, # p<0.05 compared to side, ##: p<0.01 compared to side.)

The resulting fluorescent signal localized to the nuclear envelope, consistent with a label at the nuclear lamina. We verified that the nuclear stiffness in GFP-nesprin-2/mCh-LAC cells is similar to that in GFP-nesprin-2 cells by micropipette aspiration (Supplemental Figure 2C). We also validated that the passage time through 2 μm constrictions is similar in the two cell types: the GFP-nesprin-2/mCh-LAC cells pass through the constriction on average in 140 +/−30 minutes, compared to 150+/−30 minutes for the GFP-nesprin-2 cells. Therefore, the addition of mCherry on endogenous lamin A/C does not significantly affect nuclear mechanical properties or migration times through constrictions in our assays.

GFP-nesprin-2 accumulation is confirmed in cells with the additional lamin modification (Figure 2), as well as the lower (but not statistically significant) accumulation in the 15 μm channels. Rather than accumulating at the front, mCh-LAC intensity increases at the sides of the nucleus (Figure 2A and Supplemental Figure 2E). These results were consistent with immunofluorescence results (Supplemental Figure 2F), confirming that tagged lamins behave similarly to the endogenous lamins.

Confocal imaging reveals that this increase in intensity correlates with wrinkling of the nuclear envelope at the edges of the pillars during deformation (Supplemental Figure 2G). The wrinkling at the edge of the pillars is visible in both the GFP-nesprin-2 and mCh-Lmna channels. We confirmed this wrinkling also occurs in the GFP channel in cells that do not carry the lamin modification (Supplemental Figure 2D). This wrinkling thus contributes to increasing the apparent signal at the edges of the pillars, but is an imaging artefact rather than an increase in protein accumulation at the edges of the nucleus.

Measurements of the volume and surface area deduced from the cross-sectional area and height of the micropipette and migration devices indicate that the surface area of the nuclear envelope is conserved during deformation whereas the volume decreases (Supplemental Figure 2H and I). Altogether, we find that accumulation of GFP-nesprin-2 is independent of A-type lamins.

### Nesprin-2 is moved to the front from the rest of the nuclear envelope

To determine whether the increase in nesprin levels at the front may be due to newly synthesized protein or to translocation from elsewhere at the nuclear envelope we block protein translation using cycloheximide. We find that cells still display nesprin accumulation when migrating through narrow constrictions (Supplemental Figure 3A), indicating that the accumulation is independent of protein synthesis. We estimate the time required to synthesize nesprin-2 giant by bleaching the entire nucleus in migration devices and find that after 6 hours, the nesprin signal is still weak (25% +/− 2%, N=30) compared to other cells (Supplemental Figure 3B). Therefore, the characteristic time for protein synthesis (>6 hours) is longer than for passage through the constriction (2.5 hours). Moreover, the signal is regained uniformly around the nucleus rather than exclusively at the front in deformed nuclei, indicating that the newly synthesized protein is not selectively inserted at the front of the nucleus (Supplemental Figure 3B). Protein synthesis is therefore unlikely to account for the accumulation at the front of the nucleus. Instead, nesprin must be moved along the surface of the nucleus.

**Figure 3:**
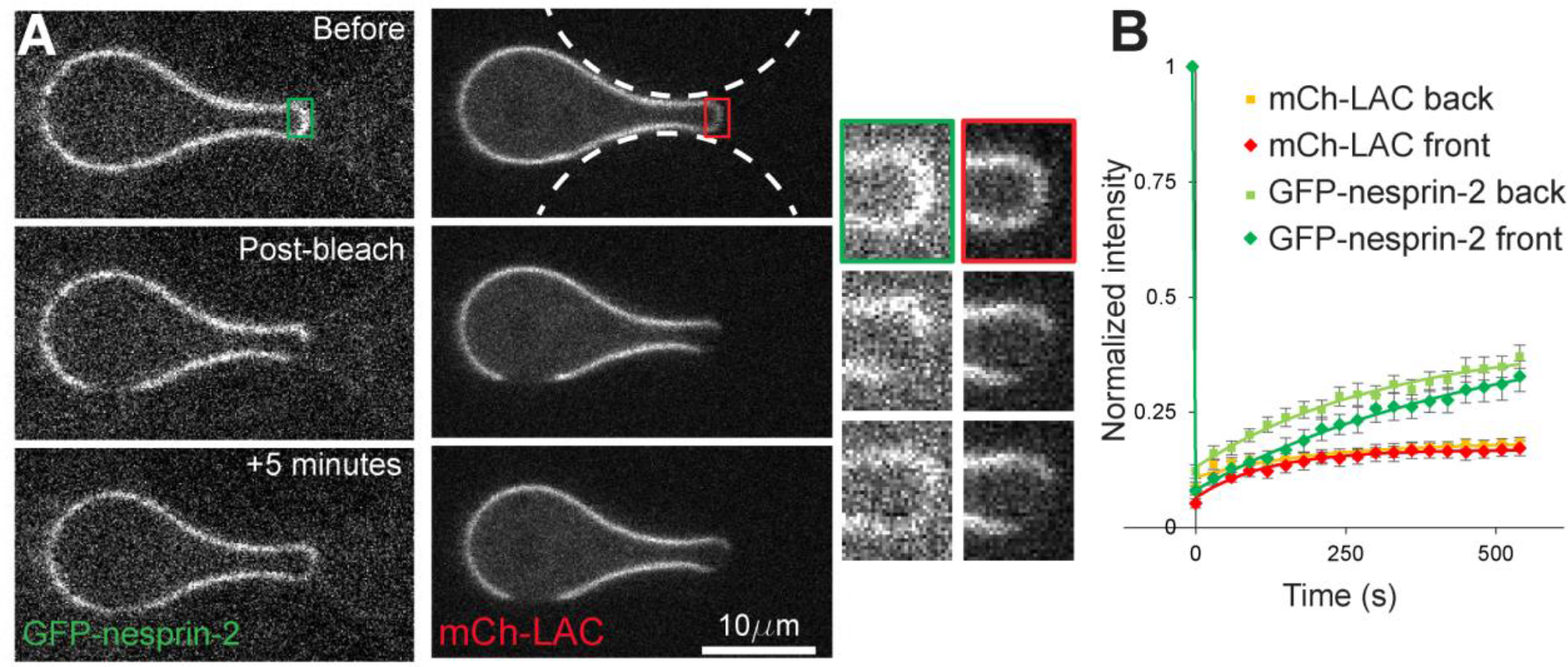
Nesprin mobility is greater than lamin A/C mobility. **A**, The deformed nucleus of cells migrating through narrow constrictions was bleached at the front and the side to determine mobility. (See also movie 3.) Inset are shown close-up views of the bleached areas at the front of the nucleus. **B**, Quantification of the recovery. (Error bars correspond to the standard error of the mean, n=24 over 3 experiments.)

To test this directly, we characterize the mobility of nesprins and lamins in our CRISPR-modified cells by bleaching two regions of the nucleus (at the front and at the back) and observing fluorescence recovery over 10 minutes (Figure 3A,B). A-type lamins demonstrate a rapid recovery of a small fraction of the fluorescence within the first 30 seconds, (approximately 15%) representing soluble lamins in the nuclear interior moving into the photobleached area, in agreement with previous results.^26^ This is followed by a plateau that represents the very slow turnover of the nuclear lamina. The recovery of nesprin-2 is gradual and is likely due to 2-D transport within the nuclear envelope as there is no soluble pool of this transmembrane protein. There is a shift in the recovery between the two regions, likely due to a higher initial fluorescence intensity at the front of the nucleus. When imaged at later time points, we find that the signal at the front is recovered within 1-2 hours after photobleaching, whereas long-term recovery is slower at the back (Supplementary Figure 3C). This more rapid recovery at the front is likely due to redistribution of nesprin-2 from the back and sides of the nucleus towards the front. The time scale of the recovery is similar to the time required to migrate through the constriction (150 minutes). We thus find that nesprin-2 is redistributed towards the front, on a time scale that is similar to the time required for its accumulation during migration through narrow constrictions.

### The actin-binding domain of nesprin-2 is necessary and sufficient to induce nesprin-2 accumulation at the front of the nucleus

To determine the role of the various domains of nesprin-2 giant we use a mini-nesprin construct that contains the actin-binding domain and only 4 of its 56 spectrin repeats (and cannot bind fascin or FHOD1), and a mutant version (I128A, I131A)^22^ that is unable to bind filamentous actin (Figure 4A).^21^ Cells expressing these constructs still show localization of endogenous nesprins at the nuclear envelope (Supplemental Figure 4A).Similarly to the GFP-nesprin-2 cells (Figure 1C, 1G, 2A, 2B), cells expressing the mini-nesprin construct show an increase in GFP intensity at the front of the nuclei as cells migrate through narrow constrictions (Figure 4A, 4C, Supplemental Figure 4B), which had not been previously described. Conversely, cells expressing the mutant construct do not show increased accumulation at the front of nuclei (Figure 4C, Supplemental Figure 4B). Furthermore, we observe that when the cell changes migration direction, the nucleus inverts its front and back, and nesprin decreases at the former front and increases at the new front (Supplemental Figure 4C, movie 6).

**Figure 4.**
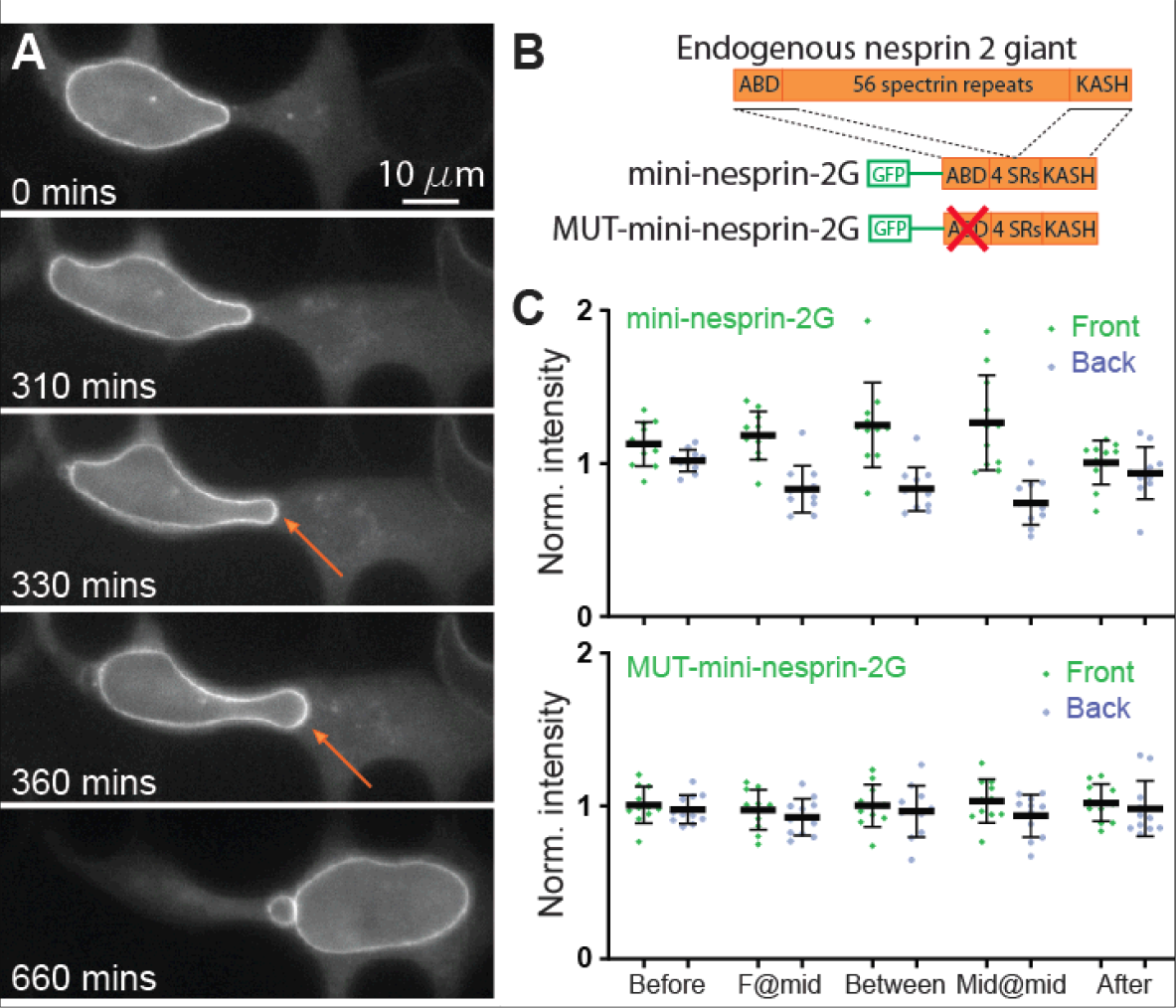
Nesprin accumulation is due to the actin cytoskeleton. **A**, Expression of the artificial mini-nesprin-2G construct results in significant nesprin accumulation in cells migrating through narrow constrictions. (See also movie 4.) **B**, Schematic depicting the endogenous protein, and the artificial mini-nesprin-2G and MUT-mini-nesprin-2G constructs. Both constructs contain the first two and last two spectrin repeats of the endogenous protein; the mutant construct presents two mutations on the actin-binding domain that prevents it from binding to actin. **C**, Quantification of the mini-nesprin-2G accumulation of cells migrating through narrow constrictions. Note that the mutant construct does not accumulate at the front of the nucleus. (See also movies 4 and 5.) (Error bars correspond to the standard deviation. Mini-nesprin-2G: n=10 cells over 4 different experiments; Mutant: n=11 cells over 4 different experiments.)

To confirm the role of the actin-nesprin link in this accumulation, cytochalasin D (1μM) is added to the cells once they are already migrating in the devices. As expected, cell migration is reduced. Half the nuclei observed back out of the constriction (N = 7 out of 13), even when the cell body remains adhered to the surface, demonstrating that actin filaments are responsible for deforming the nucleus (Supplemental Figure 4D). When the nucleus remains in the constriction, there is no accumulation of nesprin at the front (Supplemental Figure 4E). These results indicate that actin plays a determinant role in deforming the nucleus and enriching nesprin at the leading edge of the nucleus.

To characterize whether microtubules are involved in nesprin accumulation, we use a concentration of nocodazole (1 μM) that is sufficient to arrest mitosis in the migration device. Under these conditions, migration speed decreases but cells can translocate nuclei through the constrictions and we observe similar GFP-nesprin-2 accumulation as in the absence of nocodazole (Supplemental Figure 4F, H). Microtubules are thus not necessary for nesprin-2 accumulation at the front of the nucleus. Altogether, these results reveal that actin and the actin-binding domain of nesprin-2 are responsible for nesprin accumulation at the front, independently of microtubules. Furthermore, fascin and FHOD1, which cross-link actin to nesprin-2 at domains that are absent in mini-nesprins, are not necessary for nesprin-2 accumulation.

### Actin cables form a barrel around the nucleus and colocalize with nesprin-2 structures

To determine the architecture of actin around the nucleus as it is being pulled through the 2 μm constriction, we fix cells expressing GFP-nesprin-2, label them with fluorescent phalloidin and observe them on a confocal microscope. Actin is organized in cables aligned in the direction of migration around the periphery of the nucleus and it is accumulated along the edges of the pillars and at the top and the bottom of the cell surface (Figure 5A). In particular, we observe aligned structures of nesprin-2 that colocalize with actin cables along the surface of the nucleus (Figure 5B), reminiscent of TAN lines^22^ observed during cell polarization. These results are consistent with mechanical links between nesprin-2 and actin at the surface of the nucleus.

**Figure 5.**
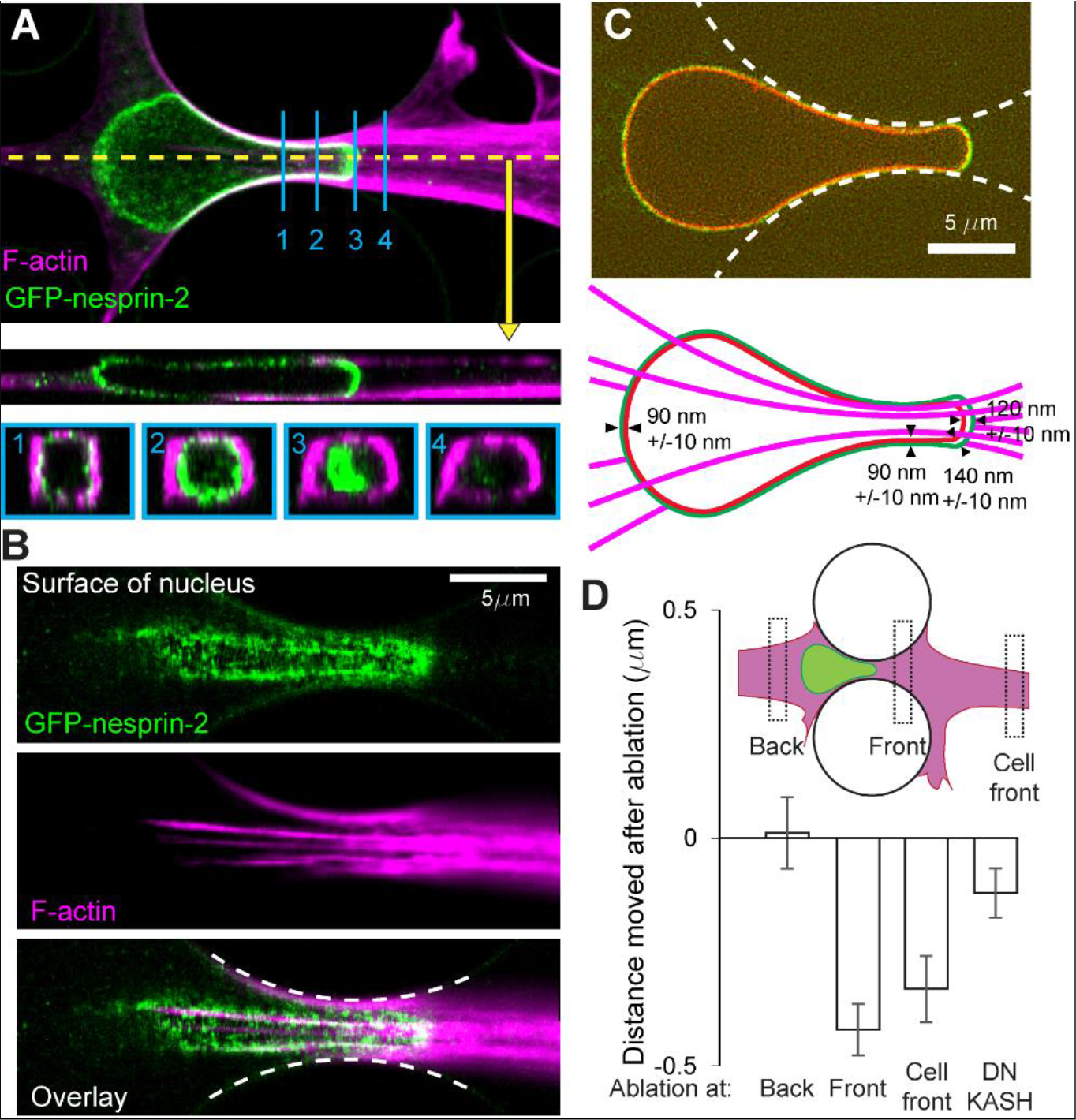
The recruitment of nesprins is due to nesprins moving to the front of the nucleus. **A**, Fixed cells labelled with phalloidin shows filamentous actin accumulation on the edges of the pillars and along the top and bottom surfaces. Cross-sections along the direction of migration (yellow) and perpendicular to the direction of migration (blue, 1-4) are shown below. **B**, A confocal slice on the surface of the nucleus shown in A indicates that GFP-nesprin-2 co-localizes with actin cables at the surface of the nucleus. **C**, Spinning disk confocal microscopy on live cells reveals the elongation state of nesprins at the nuclear envelope. The distance between mCh-LAC and GFP-nesprin-2 was measured at the back, at the constriction, at the front near the constriction, and at the center of the front. (N = 17 cells, over 3 experiments. The alignment of the two channels was corrected using at least 28 fluorescent beads.) **D**, Two-photon laser ablation reveals that ablation at the front of the nucleus results in a rearward movement of the nucleus, but not ablation at the back. The nuclei of DN-KASH, which show a decrease of endogenous nesprin at the nuclear envelope (see Supp. Fig. 4A), show a dampened rearward movement upon ablation at the front of nuclei. (Back: N=9, Front: N=30, Cell Front: N = 16, and DNKASH: N=31)

Importantly, we do not observe accumulation of actin at the front of the nucleus, only around the edges (Figure 5A, panels 3 and 4). Under our experimental conditions in live cells, although SIR-Actin absorbs onto the surface of the PDMS and creates a high background, spinning disk microscopy confirms that actin filaments are organized around the nucleus parallel to the direction of migration (Supplemental Figure 5).

The distinct markage of the actin-binding domain of nesprin, and lamin at the nuclear interior, allows for a measurement of the distance between the nuclear lamina the distal end of nesprin. As the distance between nuclear membranes is kept constant by the LINC complex,^27,28^ we expect this measurement to reflect the extension of nesprins into the cytoplasm. Measurements around the confined nucleus reveal that nesprins are more elongated at the front than elsewhere (Figure 5C, D). In particular, the elongation does not peak at the center of the front (122 nm) but is higher close to the constriction (140 nm), in spite of very low values in the constriction (85 nm). This is consistent with our finding that actin is organized around the periphery of the nucleus and the cell, but not at the front of the nucleus, where we observe accumulation of GFP-nesprin-2. Altogether these results suggest actin pulls on nesprin-2 at the periphery of the nucleus.

### Actomyosin contractility and linear actin nucleation are necessary for migration through constrictions

To determine the actin architectures responsible for translocating the nucleus through constrictions, we used a panel of drugs that disrupt branched actin nucleation (CK666, an Arp2/3 complex inhibitor), linear actin nucleation (SMIFH2, inhibits the FH2 domain of formins) and that inhibit myosin II (para-nitroblebbistatin) and ROCK (Y-27632, upstream of myosin). We treated cells in the migration devices and observed the time required to migrate through narrow constrictions and the same distance in wide control channels (as described previously)^4^. We found that the time required to migrate through the narrow constrictions was increased in the cells treated with formin, myosin and ROCK inhibitor (Supplemental Figure 5b). Only one cell treated with ROCK inhibitor migrated through the constrictions within the time frame of the experiment. Migration times in the control channel were increased in cells treated with the Arp2/3 complex inhibitor and decreased in cells treated with ROCK inhibitor.

### Forces are applied to deforming nuclei at the front of the nucleus

We performed ablation experiments to determine whether the nucleus is pulled from the front during migration through narrow constrictions, or whether contraction at the back of the cell is responsible for pushing the nucleus through. We ablated material 5 μm in front or behind cells migrating through narrow constrictions. Ablation at the front resulted in an immediate displacement of the nucleus towards the back, which we did not observe when the nucleus was ablated at the back (Figure 5D). We confirmed these results by ablating material further at the front of the cell and observed a similar rearward displacement of the nucleus. In cells expressing DN-KASH, in which nesprins are displaced from the nuclear envelope, ablation at the front of the nucleus resulted in significantly reduced rearward displacement.

## DISCUSSION

We show here that giant nesprin-2 accumulates at the front of the nucleus during migration through narrow constrictions. This accumulation depends on the actin-binding domain of nesprin-2 and does not depend on microtubules or A-type lamins.

### Labelling the actin-binding domain of nesprin-2 tags the giant isoform

Here we tag endogenous nesprin-2 by inserting a GFP sequence at the N-terminus of the gene, immediately before the sequence that encodes the actin-binding domain. Whereas other nesprin-2 isoforms have been described^8^ and observed^29^ only the giant isoform contains both the actin-binding domain and the KASH domain, which determines nesprin localization at the nuclear envelope. The localization of GFP fluorescence at the nuclear envelope shows we are specifically labelling the giant isoform.

### The nuclear envelope does not stretch when constricted, but instead folds

Previous studies showed nuclear envelope folding as cells migrate through narrow constrictions.^4,23^ It was unclear whether this behavior was an artefact caused by increased nuclear envelope stiffness due to lamin A/C overexpression in these cells (used to label the nuclear envelope). Here we confirm that the nuclear envelope folds in cells expressing wild-type levels of lamins and in which the stiffness of the nucleus has not been altered, as demonstrated by micropipette and migration experiments. Nuclear envelope stretching at the poles could explain nuclear envelope rupture at the front and back of the nucleus observed during migration through narrow constrictions.^30,31^ In our measurements we did not observe any indication that the nuclear envelope is stretched during deformation. Instead, we found that the volume decreases and the surface area of the nucleus remains constant.

### Nesprin-2 moves to the front of the nucleus

We show that nesprin accumulation at the front of deforming nuclei is not due to newly synthesized protein. Nesprin-2 accumulation at the front of the nucleus is thus likely due to the movement of nesprins from elsewhere on the nucleus to the front. This accumulation could be due to active transport to the front of the nucleus by the actin cytoskeleton, or due to passive diffusion and retention at the front of the nucleus (or both). In the latter case, a (not yet identified) ligand of nesprin-2 at the front of the nucleus would be required to retain it. In both cases, the longer it takes to translocate the nucleus through the constriction, the more nesprin will be found at the front of the nucleus. Indeed, in support of these mechanisms, we find there is a positive correlation between the time required to translocate the nucleus through the constriction and the nesprin-2 accumulation (Supplemental Figure 2F). We considered altering the mobility of nesprins at the nuclear envelope by targeting the proteins (lamin, SUN) that link nesprins to the inner nuclear membrane to determine whether the accumulation could be due to passive diffusion. However, this is likely to alter nuclear mechanics and could result in displacement of nesprins to the ER. We therefore find that nesprin-2 moves to the front of the nucleus either actively, or by retention by an unidentified ligand.

### Nesprin accumulation is due to the actin cytoskeleton

We studied the role of actin and microtubules, two cytoskeletal filaments that associate with motors to exert forces within the cell. We find that the accumulation is associated with actin, but not with microtubules. Moreover, accumulation is selectively dependent on the actin-binding domain, and not on known crosslinkers of actin and nesprin-2 (fascin and FHOD1).

We find that actin filaments are organized in a barrel around the nucleus parallel to the direction of migration. We identify linear nesprin-2 structures that associate with actin filaments at the surface of the nucleus. Our results are consistent with previous observations in which non-muscle myosin IIb accumulates on filamentous structures (presumably actin) around the nucleus as it translocates through constrictions.^17^ Endogenous nesprin-2 may thus bind to filamentous actin at the nuclear periphery, and could be pulled to the front of the nucleus during displacement of actin filaments, or could slide to the front along the actin cables.

### The nucleus is pulled through narrow constrictions

Our experiments demonstrate that there is an elastic recoil when material is ablated at the front of the nucleus, consistent with a pulling force from the front of the nucleus. This is in agreement with the description of nuclear movement in 2-D,^32^ the nuclear piston,^20^ and predictions based on the patterns of strain in the nuclear interior.^19^ Moreover we demonstrate that inhibiting formin and myosins reduces the efficiency of nuclear translocation through constrictions, consistent with a mechanism that relies on linear actin nucleation and actomyosin contractility. We have observed that when adhesion at the front of the cell is lost, the nucleus of the cell falls back (movie 7). All of these results point to a mechanism in which the nucleus is pulled through the constriction via actomyosin contractility at the front of the cell. The sharp decrease in recoil distance in DN-KASH cells points to the involvement of nesprins in this process. Our results thus indicate a pulling force that is LINC complex-dependent.

This data does not preclude additional mechanisms that could contribute to translocating the nucleus through constrictions. Indeed, contraction of the cell rear may be sufficient to translocate DN-KASH cells. Nevertheless, it is not sufficient to maintain nuclear deformation when adhesion is lost at the front of the cell in non-DN-KASH cells, as demonstrated in ablation experiments (Figure 5) and movie 7. We therefore find that the dominant mechanism is a pulling force exerted at the front.

## OUTLOOK

We show here that the link between actin and nesprin drives nesprin-2 giant accumulation at the front of nuclei during cell deformation through narrow constrictions. More work is required to determine whether nesprin-2 is involved in pulling the nucleus forward through constrictions. The organization of actin around the nucleus and the elongation of nesprin-2 giant indicates that the forward pull occurs at the sides of the nucleus rather than at the front. This mechanism could allow the force on the nucleus to be distributed over a wider area, limiting the stress and damage that could occur if the force was concentrated at a small area at the front of the nuclear envelope.

## ACKNOWLEDGEMENTS

Antibodies were generously provided by G. G. Gundersen and the Wolfson center for inherited disease. Reagents were generously provided by Kristine Schauer (Institut Curie), Marius Doring (Institut Curie) and Michel Wassef (Institut Curie). This work was carried out in part at the Cell and Tissue Imaging (PICT-IBiSA) and Nikon Imaging Centre, Institut Curie, member of the French National Research Infrastructure France-BioImaging [ANR10-INBS-04]. This work was supported by La Ligue contre le Cancer [REMX17751 to PMD], the Agence Nationale de la Recherche [ANR Actin2Nucleus to TB, CS, NB, BC] and Fondation ARC [PDF20161205227 to PMD].

The authors declare that they have no conflict of interest.

## STAR METHODS

### KEY RESOURCES TABLE

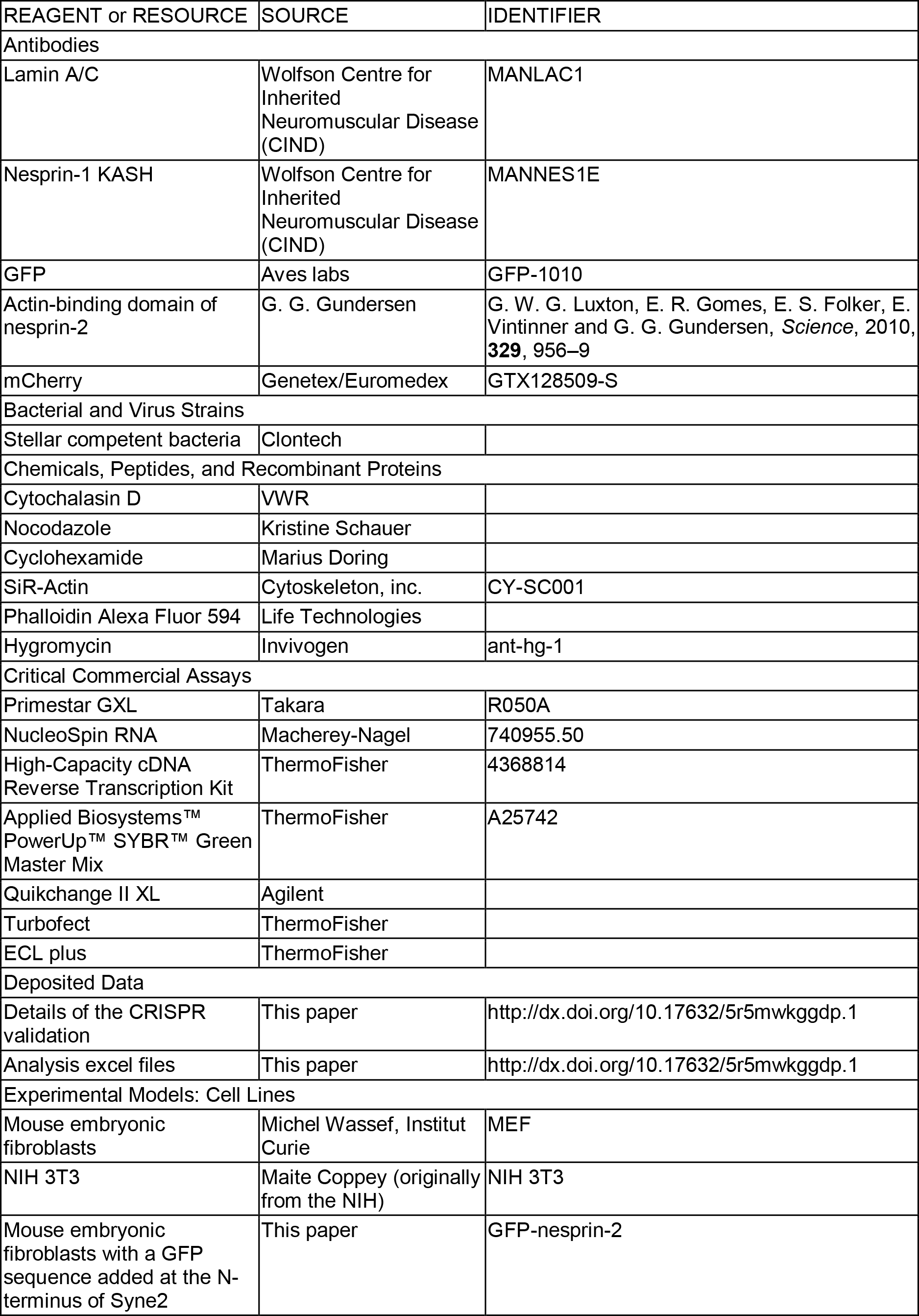
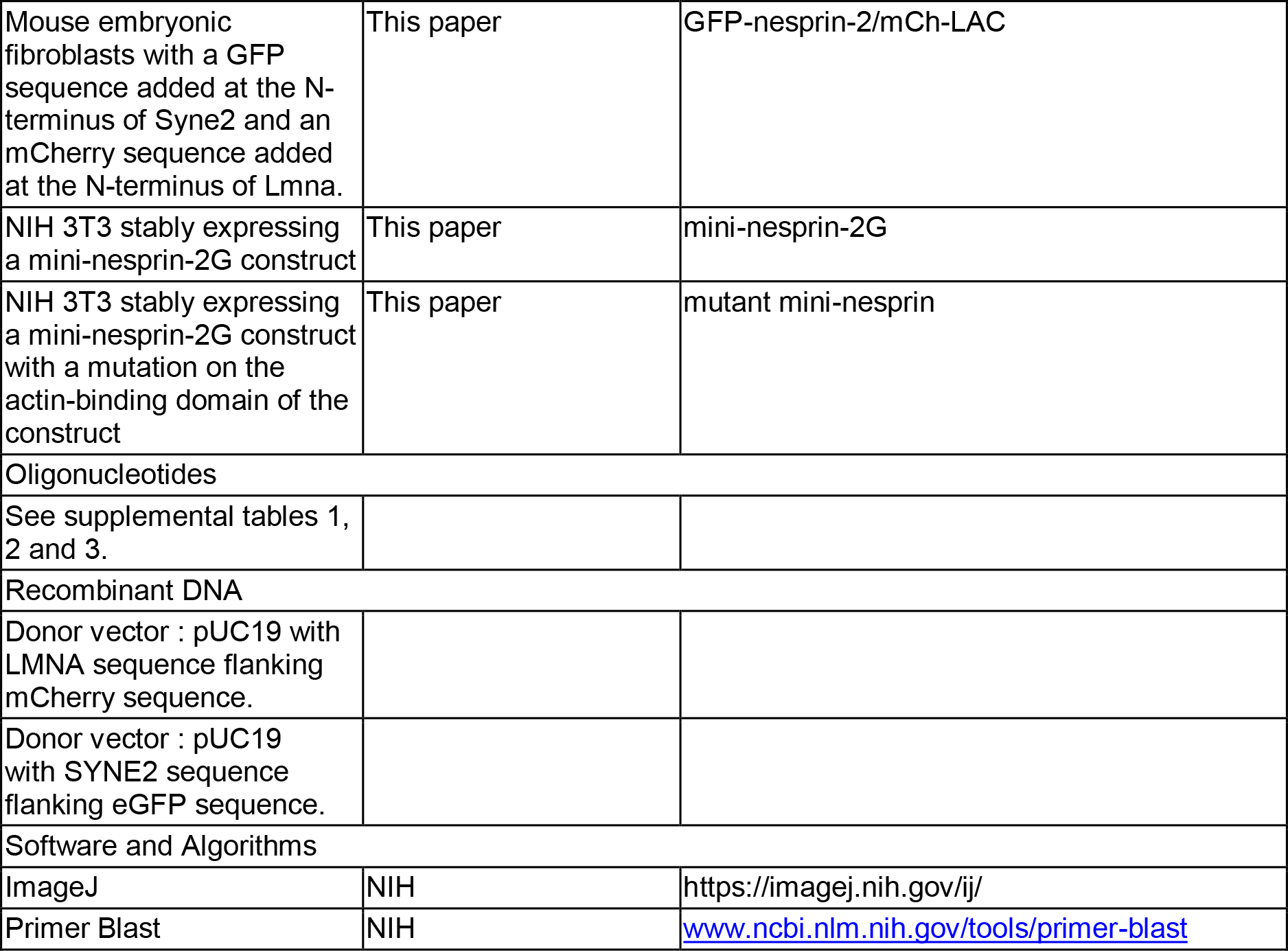

### CONTACT FOR REAGENT AND RESOURCE SHARING

Further information and requests for resources and reagents should be directed to and will be fulfilled by the Lead Contact, Patricia Davidson

## EXPERIMENTAL MODEL AND SUBJECT DETAILS

### Mouse embryonic fibroblasts

Mouse Embryonic Fibroblasts (MEF) were obtained from M. Wassef (Institut Curie). NIH3T3 MEFs were obtained from M. Coppey (Institut Jacques Monod) They were cultured at 37 degrees Celsius in a humidified incubator with 5% CO_2_. The cells were cultured in DMEM (Life Technologies) supplemented with 10% (v/v) Fetal Bovine Serum (Hyclone) and 1% Pennicilin/Streptomycin (Life Technologies). For imaging experiments in the microfluidic migration devices, the medium used consisted of DMEM without phenol red and with HEPES (15 mM) (Life Technologies), supplemented with 10% Fetal Bovine Serum (Hyclone), 100 units/mL Penicillin, and 100 μg/mL Streptomycin (Life Technologies).

## METHOD DETAILS

### Reagents

Phalloidin Alexa Fluor 594 (Life Technologies) was used at a dilution of 1:50 to label filamentous actin. For western blotting experiments, we used antibodies against mCherry (GeneTex, 1:2000), Lamin A/C (MANLAC1, Wolfson Centre for Inherited Neuromuscular Disease, 1:100),^33^ GFP (Aves labs, 1:2000), the actin-binding domain of nesprin-2s^22^ generously provided by Greg G. Gundersen (1:3000), and actin (Santa Cruz,1:250). For immunofluorescence experiments we used an antibody against the KASH domain of nesprin-1 (MANNES1E, Wolfson Centre for Inherited Neuromuscular Disease, 1:100)^34^.

Cytochalasin D (VWR) was dissolved to a stock concentration of 5 mg/ml (10 mM) in dry DMSO and used at a working concentration of 0.5 μg/ml (1 μM). Nocodazole at a stock concentration of 1mM in DMSO was graciously provided by Kristine Schauer (Institut Curie) and used at a working concentration of 1 μM. Cyclohexamide at a stock concentration of 100 mg/ml in DMSO was graciously provided by Marius Doring (Institut Curie) and used at a working concentration of 10 μg/ml. SiR-Actin (Cytoskeleton, Inc.) at a stock concentration of 1 mM in DMSO was diluted to 250 nM in imaging medium and added to the cells 4-6 hours before seeding into the microfluidic devices. SMIFH2 (VWR) was dissolved to a stock concentration of 50 mM in dry DMSO and used at a working concentration of 50 μM. CK666 (VWR) was dissolved to a stock concentration of 250 mM in dry DMSO and used at a working concentration of 0.25 mM. Para-nitrobelbbistatin (Axol) was dissolved to a stock concentration of 37 mM in dry DMSO and used at a working concentration of 37 μM. Y-27632 (VWR) was dissolved to an aqueous stock concentration of 100 uM and used at a working concentration of 1 μM.

### Creation of two CRISPR-modified cell lines

Syne2 modification: we inserted the DNA sequence for green fluorescent protein (GFP) and a flexible linker sequence immediately after the start codon (ATG) of *Syne2*. Lmna modification: we inserted the gene sequence for mCherry (mCh) and a flexible linker immediately after the start codon (ATG) of *Lmna*. The sequences were designed based on strategies detailed in Wassef et al.^35^

### DNA cloning

CRISPR single guide RNAs (sgRNA) were designed using the Optimized CRISPR Design tool (http://crispor.tefor.net/). Sequences are provided in Supplementary Table 2. For sgRNA-encoding plasmids, single-stranded oligonucleotides (Eurofinsgenomics.eu) containing the guide sequence of the sgRNAs were annealed, phosphorylated and ligated into BbsI site in px335 (kindly provided by M.Wassef, Curie Institute, Paris, France). To construct the donor vector, homology arms of approximately 800 base pairs (bp) were amplified from genomic DNA using PCR primers. Primers were designed with 40 bp overhangs compatible with a pUC19 backbone (see supplemental table 1 and 2). The plasmid was digested with Xba1 and Ecor1 (New England Biolabs). mCherry was amplified with a 20 bp overhang, and the linker sequence SGLRSRAQAS was added in the right arm forward primer (sequences are provided in Supplementary Table 2). Gibson reactions were performed using a standard protocol with a home-made enzyme mix.^36^ See Supplemental Tables 1 and 2 for cloning primers.

### Clonal selection of cells and PCR validation

Ten million mouse embryonic fibroblasts (ATCC) were transfected with 90uL of Polyethylenimine (PEI MAX #24765 Polysciences) diluted in 240uL NaCl 150mM and 15ug of the pX335-gRNA and pUC19-homology arms-mCherry plasmids also diluted in 240uL NaCl 150mM. Seven days after transfection, fluorescent cells were selected on a BD FACS ARIA II. After a further ten days, cells are sorted again, individual fluorescent cells are placed in individual wells in a 96 wells plate. Once they have reached confluence, the clonal populations are split into two plates, one for PCR screening and the other for backup. To perform the PCR screening, genomic DNA was isolated from each well of a confluent 96-well plate according to previous protocols.^35^ An aliquot of 1 μL of this lysate was used in a 25 μl PCR reaction. PCR reactions were performed using Primestar GXL Takara, according to manufacturer’s instructions. For each targeted locus, two sets of genotyping primers spanning the junction of genomic sequences and targeting vector were used (left and right arms). Gene-specific primers were designed outside the 5’ and 3’ homology arms. To identify insertions or deletions due to errors during non-homologous end joining, the region flanking the sgRNA target sites (100 bp) was amplified using PCR with gene-specific primers and directly assessed by Sanger sequencing (Supp. Fig. 1). Sequences of all primers are provided (see Supplementary Tables 1 and 2). See Supplemental Tables 1 and 2 for PCR primers.

### Validation of clonal populations

Several clonal populations were picked and cDNA from bands obtained in the three PCRs was sequenced to check for mutations. From these results we selected a clonal population of MEFs with a homozygous knock-in of the GFP gene on Syne2 devoid of mutations in the targeted region. Similarly, several heterozygous clonal populations of cells with an mCherry knock-in on the Lmna gene were sequenced. We thus obtained 6 clonal populations without mutations and verified that the addition of the extragenous sequence did not affect the mechanical properties of the cells. Following mechanical validation, we picked a heterozygous cell line with similar properties to the parental cell line for further experiments.

### Validation of protein by Western Blot

In both cell lines obtained, lysates of the cells were run on a western blot and probed with antibodies for the endogenous protein and fluorescent marker added (Supplemental Figure 1). The migration was performed in Tris-Acetate gels (Thermo) to reveal giant nesprins or polyacrylamide gradient gel to reveal lamin A/C (made in house). We verified the localization of a GFP band that corresponded to a band detected using a Nesprin-2 antibody. After picking the mCh-LAC/GFP-nesprin-2 clone that had the most similar mechanical properties, we verified the presence of bands 22 kDa higher than the endogenous lamin A/C bands, consistent with the size of the mCherry protein. These higher bands are also revealed by an mCherry antibody (Supplementary Figure 1). We determined that the proteins carrying the mCherry tag are less abundant than the non-tagged proteins: the combined intensity of the lamin A/C bands around 90 kDa is approximately 10% of the intensity of the bands at 70 kDa.

### Quantitative PCR

RNA was extracted with a NucleoSpin RNA kit (Macherey-Nagel, 740955.50) following manufacturer’s instructions. cDNA was synthetized from 2ug of RNA using a multiscribe High capacity cDNA reverse transcription kit (ThermoFisher 4368814). The primers were designed using Primer blast software (www.ncbi.nlm.nih.gov/tools/primer-blast).

mRNA expression was performed using the Applied Biosystems PowerUp SYBR Green Master Mix (ThermoFisher A25742). qPCR reactions were run on a steponeplus (ThermoFisher). The data was analysed using the Delta Ct method and using tata-binding protein as a control. See Supplemental Table 3 for qPCR primers.

### Creation of stable NIH 3T3 mini nesprin cell lines

Plasmids were cloned into a pcDNA3.1 hygro(−) vector (ThermoFisher), digested by BamHI using a In-fusion HD cloning kit. The cytoplasmic and KASH domains of the mini-nesprin constructs were obtained from a previous publication.^22^ The mutant domain was made from this construct by site-directed mutagenesis (Quikchange II XL, Agilent). The DN-KASH construct was made by ligation after digestion by ClaI of a PCR product from the mini-nesprin construct. For sequences see Supplementary Table 4. Constructs were verified by sequencing coding regions.
NIH 3T3 cells were transfected plasmids Turbofect according to manufacturer’s instructions. (See Supplemental Table 4 for plasmid sequences.) The cells were cultured for two weeks in medium supplemented with 200 μg/ml hygromycin and sorted by fluorescence-activated cell sorting (FACS) based on the green signal of the GFP.

### Western Blotting

Protein lysates were prepared in RIPA buffer (20 mM Tris HCl pH 7.4, 150 mM NaCl, 1 mM EDTA pH 9, 0.1% SDS, 1% Triton-X 100). Cells were grown to confluence in a 10 cm petri dish. The cell monolayer was rinsed with Phosphate Buffered Saline and 500 μL RIPA buffer containing protease inhibitor (Sigma) was added to the dish. The cell layer was scraped and the solution was transferred to a cold Eppendorf tube on ice. After 20 minutes of incubation, the tube was centrifuged at 10000g for 10 minutes and the supernatant was transferred to a new tube with sample buffer, for a final concentration of 0.0675 M Tris, 10% (v/v) glycerol, 20 g/l SDS, 1% (v/v) 2-mercaptoethanol and 5 mg/l bromophenol blue. A small aliquot was kept to assess total protein content using a BCA reagent assay (Thermo Scientific). Acrylamide gels were prepared in-house with a concentration gradient of 4-14%. Appropriate volumes of the protein solutions were loaded on the gels and they were run in SDS-page buffer (0.25M Tris base, 2M glycine, 1% (wt/wt) SDS) at 200V for 10 minutes, then 170V on ice for 1 hour. The proteins were then transferred to a methanol-activated PVDF membrane in sample buffer supplemented with 20% (v/v) ethanol at 16V for 20 hours. After transfer the membrane was soaked in 2% (w/v) milk in Tris-Buffered Saline containing 0.5% (v/v) Tween (TBS-T) for 20 minutes. Primary antibodies were incubated for 2 hours at room temperature and in Secondary antibodies for 1 hour at room temperature. Antibodies were diluted in 2% milk solution in TBS-T. Bands were revealed with ECL Plus reagent (ThermoFisher) on an Amersham AI680 (G.E.) and analysed with ImageJ/FIJI software.

### Microfluidic micropipette aspiration

Micropipette aspiration devices were prepared as detailed previously.^37^ Timelapse imaging was performed on an inverted Nikon Ti-E equipped with a sCMOS camera (2048 ORCA Flash4.0 V2, Hamamatsu), a 60x objective (Nikon), and a temperature control chamber (Tokai). Cells solutions were prepared by passaging cells, centrifuging to remove the medium and suspending the pelleted cells at 5 × 10^6^ cells/mL in a solution containing 20 g/L bovine serum albumin (Sigma), 0.2% (v/v) fetal bovine serum (Hyclone) and 10 μg/mL Hoechst 33342. They were kept on ice until the measurements. Cells were perfused into the device and brightfield and fluorescence images were taken every five seconds. Analysis of the data was performed using ImageJ/FIJI to determine the protrusion length as a function of time.

### Migration devices

The microfabricated migration devices were prepared as described previously.^4,23^ Timelapse imaging was performed on the same Nikon Ti-E as above. Images of the migration devices were taken every ten minutes. The resulting images were turned into movies using ImageJ/FIJI and cells that translocated their nuclei through 2 μm constrictions were selected for analysis (both in the direction of net migration and across the channels, see Supplementary Figure 1E). The mini-nesprin-2 and mutant mini-nesprin cell lines displayed high mortality in the devices. For this reason, we analyzed cells migrating through 3 μm constrictions in addition to the 2 μm constrictions studied with the GFP-nesprin-2 cells.

The migration time through the constriction was defined as the time between the “Front@mid” and “Mid@mid” time points.

### Measurement of the intensity around the periphery of the nucleus

Images were analyzed using ImageJ/FIJI software. The “segmented line” tool is used to draw a line 5 pixels wide clockwise along the periphery of the nucleus, starting at the front (relative to the direction of migration). The “plot profile” tool is then used to obtain the value of the intensity along the profile and imported into excel. The profile is then split into four segments as follows: front (first and last 5% of the profile), right (5-45%), back (45-55%) and left (55-95%) (see Supplementary Figure 2A). The average value of the intensity is obtained for each segment. The background fluorescence is obtained by measuring the fluorescence in the region of a pillar and the fluorescence at the center of the nucleus is measured to evaluate bleaching due to imaging. The value of the background fluorescence is subtracted from all of the values of fluorescence obtained. The resulting intensity values are then normalized in two ways. We either divide the fluorescence values at the periphery of the nucleus by the value of the fluorescence at the center of the nucleus. In this manner we eliminate the effect of bleaching, but are subject to artefacts if the stretching of the membrane results in a decrease of the fluorescence per unit area as the nucleus goes through the constriction. This type of normalization was used to obtain the bar graphs in Supplementary figure 2B. Alternatively, we normalize the values at the front and back by dividing by the average value at the sides (average of left and right). We report this data as scatter plots obtained using PRISM software (Figure 1F, 2B, 4C).

### Fluorescence recovery after photobleaching (FRAP) experiments

Whole nucleus bleaching experiments were performed on an inverted Nikon Ti-E A1R confocal microscope equipped with a 60× objective. The GFP signal was bleached by zooming in on a nucleus and using 10 loops of the 403 nm laser. The entire area was imaged before and after bleaching to verify that the fluorescent signal of the nucleus had disappeared. The same area was imaged 6 hours later to assess fluorescence recovery (and thus protein turnover).

The quantification of fluorescence intensity was performed similarly to the analysis above. Briefly, ImageJ tools were used to draw a line along the periphery of the nucleus and obtain an average value of the intensity. The signal intensity in the region of pillars was also measured to obtain a value of the background noise. To eliminate changes due to the imaging conditions and bleaching at the two time points, unbleached nuclei were also quantified on the same image at both time points. To the best of our knowledge the same nucleus was imaged at both time points. The average of the background-subtracted intensity of the bleached nuclei was thus obtained and the average intensity of the unbleached nuclei was used as a reference.

FRAP studies on small regions of the nucleus were performed on an inverted confocal spinning disk microscope (Roper/Nikon) equipped with a FRAP module, a thermostatic chamber, lasers at 405/491/532/561nm, and an EMCCD Evolve camera. The mobility of the GFP-syne2 and mCh-LAC proteins was assessed by bleaching a region in the periphery of the nucleus at the front and the side. The bleaching was achieved by scanning the selected region over 50 loops with a 405 and 491 nm laser at 100% laser power. The area was imaged before bleaching.

The signal in the bleached area was quantified using ImageJ/FIJI software. To prevent artefacts due to bleaching of the signal during imaging, we analyzed the signal intensity in an area of the nuclear periphery that was not intentionally bleached with the laser. We also measured the background fluorescence intensity in the region of a pillar. This background value is subtracted from the intensity of the bleached area and the reference area. The normalized signal is then obtained from the ratio of the intensity of the bleached area to the reference area for each time point. The values are further normalized by setting the value of the signal before bleaching as 1. The values at all time points are thus divided by the value before the bleaching. The curves were fit using the following equation: *I* = *I*_0_ + *A*(1 − *e*^−t^/*τ*), where *I*_0_ is the initial value of the intensity, A is a constant that is used to obtain the mobile fraction and τ is the half-time.

### Volume and surface area measurements

The volume and surface area of the nuclei were approximated from values of the cross-sectional area and perimeter of the nuclei deforming through 2 μm constrictions. Epifluorescence images of the mCh-LAC channel were chosen as the signal in this channel has a higher signal-to-noise ratio, resulting in an easily segmented background. To avoid artefacts due to increased signal at the constriction, the outline of the nucleus was drawn by hand using the paintbrush tool in ImageJ software (NIH, imagej.nih.gov). The area (A) and perimeter (P) of the drawn shape were computed using the “Analyze Particles” tool in ImageJ. The volume (V) and surface area (SA) were then determined using the following equations and knowing the height (H) of the microchannels: = *A* × *H*; *SA* = 2 × *A* + *P* × *H*. The volume and surface area for each nucleus was determined and the average values are reported in Supplementary Figure 2D and E.

### Measurements of the distance between the mCh-LAC signal and the GFP-nesprin-2 signal

Stacks of images of live cells were acquired on an inverted spinning disk confocal microscope equipped with a live SR module equipped, an sCMOS camera (Flash4 Hamamatsu) and a thermostatic chamber. Images were acquired using a 100× objective (resulting a in pixel size of 65 nm) and a quad filter to allow acquisition of two wavelengths without changing filters. The nuclei were placed at the center of the field of view to minimize aberrations due to the objective. To further estimate corrections between the positions of the signal of the two colors, fluorescent beads were imaged under the same conditions. The average distance between beads (n>28) in four directions (0°, 45°, −45° and 90°) at the center of the image was used to correct the values measured.

### Immunofluorescence

Cells were plated on glass coverslips for 24 hours and fixed using 4% (w/v) paraformaldehyde. They were then rinsed with PBS and permeabilized with 0.1% (v/v) Triton X-100. After blocking with 2% bovine serum albumin (Sigma) in PBS, the cells were incubated in primary antibody for two hours at room temperature. After rinsing with PBS, the samples were incubated in secondary antibody for one hour, rinsed, mounted in 1:1 glycerol/PBS and imaged on a spinning disk microscope.

### Photoablation

Ablation experiments were performed on an Inverted Laser Scanning Confocal Microscope with Spectral Detection and Multi-photon Laser (LSM880NLO/Mai Tai Laser - Zeiss/Spectra Physics) used for cell ablation. Ablation was performed by bleaching a zone 5 μm wide with the laser set to 800 nm and a laser power of 10 %. When ablating at the “front” or the “back” of the nucleus, the ablation zone was 5 μm from the nucleus. When ablating the “cell front” each protrusion at the leading edge of the cell was ablated. The cell was imaged immediately before and immediately after the ablation and the distance by which the nucleus had moved was quantified using ImageJ.

## QUANTIFICATION AND STATISTICAL ANALYSIS

Excel (Microsoft) was used for statistical analysis. The number of cells analysed and the number of individual experiments are indicated in the appropriate figure legends.

### Analysis of the intensity around the nucleus

GFP-nesprin-2: Constriction: n=17 cells over 4 different experiments, 15 μm: n=20 cells over 3 experiments. GFP-nesprin-2/mCh-LAC: Constriction: n=19 cells over 5 different experiments, 15 μm: n=27 over 6 experiments. Mini-nesprin-2G: n=10 cells over 4 different experiments; Mutant: n=11 cells over 4 different experiments. Cycloheximide: n=6 cells over 2 experiments; Nocodazole: n=6 cells over 2 experiments. *FRAP experiments*: n=16 over 3 experiments. *6 hour bleach experiment*: n=30 cells over 2 experiments. *Quantitative PCR*: Experiments performed in triplicate on three separate lysates (obtained three separate times). *Micropipette experiments*: GFP-nesprin-2: n=76 cells over 4 experiments, GFP-nesprin-2/mCh-LAC: n=63 cells over 3 experiments. *Volume and surface area measurements:* n=15 cells over 4 experiments. *Measurements of the distance between mCh-LAC and GFP-nesprin-2*: n=17 cells over three experiments and 28 beads.

## DATA AND CODE AVAILABILITY

All excel sheets used to analyze the data and the data used to validate the CRISPR modification is available online at http://dx.doi.org/10.17632/5r5mwkggdp.1.

## Supplemental Figures

**Supplementary Figure 1:**
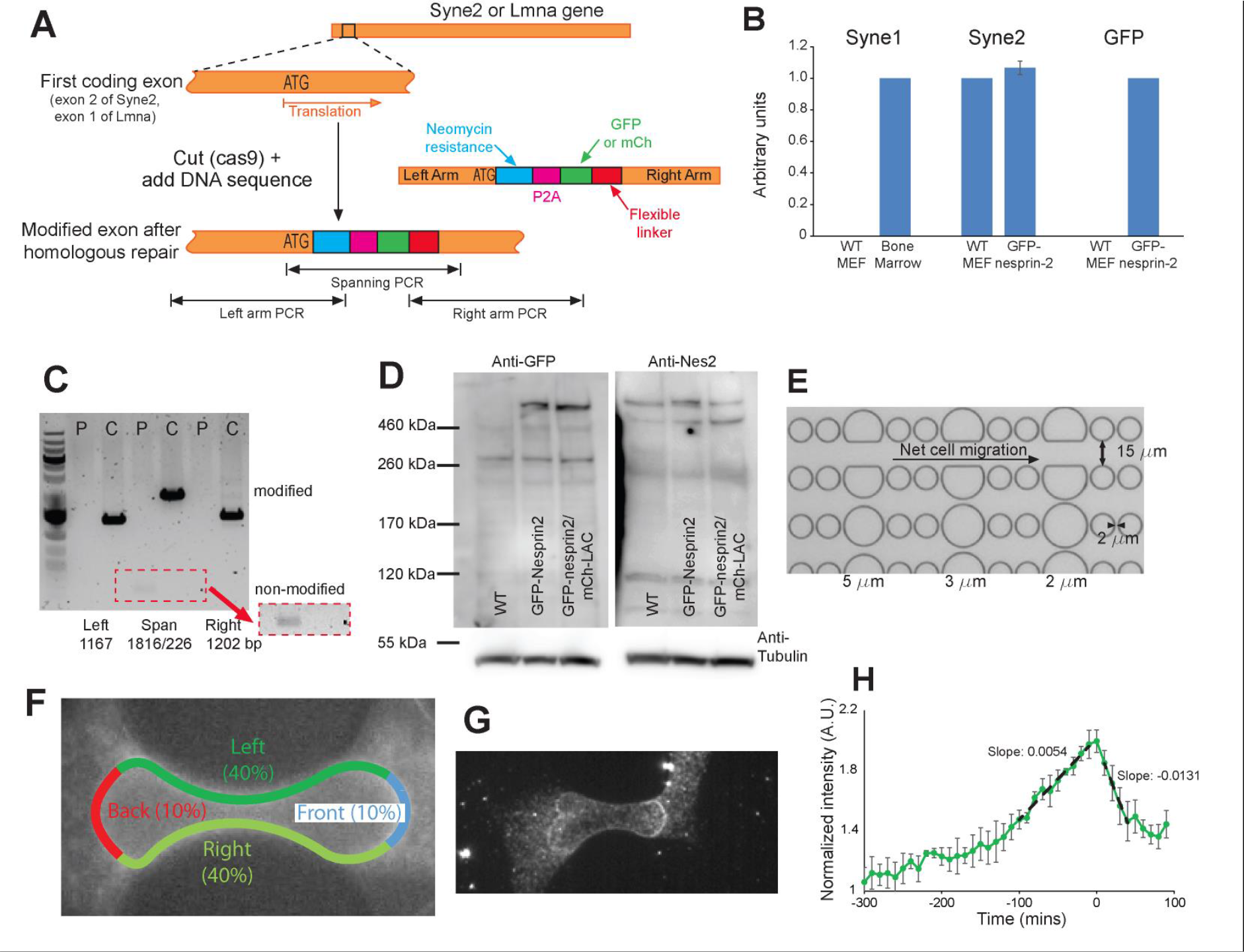
CRISPR modification and validation. **A**, Schematic showing the insertion of the CRISPR modification immediately after the start codon. A cut is induced immediately after the ATG sequence (in the second exon of Syne2 and the first exon of Syne1) using the CRISPR-cas9 system in the presence of a plasmid. This plasmid consists of the left arm (before and including the ATG), a neomycin resistance sequence, a P2A sequence, the sequence specific to the fluorescent reporter, a flexible linker and the right arm which matches the sequence of the gene immediately after the ATG sequence. Homologous DNA repair occurs in a fraction of the cells using the transfected plasmid and will insert the desired sequences into the genome. At the bottom of the schematic we show the three regions targeted by the PCR reactions. **B**, Quantitative PCR measurements. We compared the expression of the actin-binding domain of Syne1 in wild-type MEFs to Bone marrow cells, the expression of the actin-binding domain of Syne 2 in GFP-nesprin-2 cells to wild-type MEFs and the expression of the GFP-nesprin construct in GFP-nesprin-2 cells to wild-type MEFs. Error bars are included on the wild-type MEFs in Syne1 and GFP but are not visible due to the very small values. (N= 3, 4, and 2 respectively. Experiments carried out from lysates obtained on two separate days.) **C**, PCR to validate the insertion GFP-nesprin-2. The gene sequences tested are shown in panel A. The sizes expected are labelled below the image. The area outlined with a dashed red line was reproduced at higher contrast to see the band corresponding to the non-modified sequence. (P: parental cell line, C: CRISPR cell line.) **D**, Western blotting to validate the localization of a band labelled with a GFP antibody that is at the same size as bands labelled with an antibody for nesprin-2. Although two bands are expected due to the addition of a GFP to the nesprin-2, the size of the GFP (27 kDa) is small compared to the size of nesprin-2 giant (800 kDa) and bands are not resolved at such high molecular weights. **E**, Brightfield image of the migration devices. **F**, Schematic description of the regions around the nucleus analyzed. **G**, Immunofluorescence image of wild-type cells migrating through a constriction labelled with nesprin-2 actin-binding domain antibody. **H**, Quantification of the intensity at the front of the nucleus (normalized to the sides) with time. Here we averaged the data from three cells. The time at which the center of the nucleus passes through the center of the constriction is defined as time zero.

**Supplemental Figure 2:**
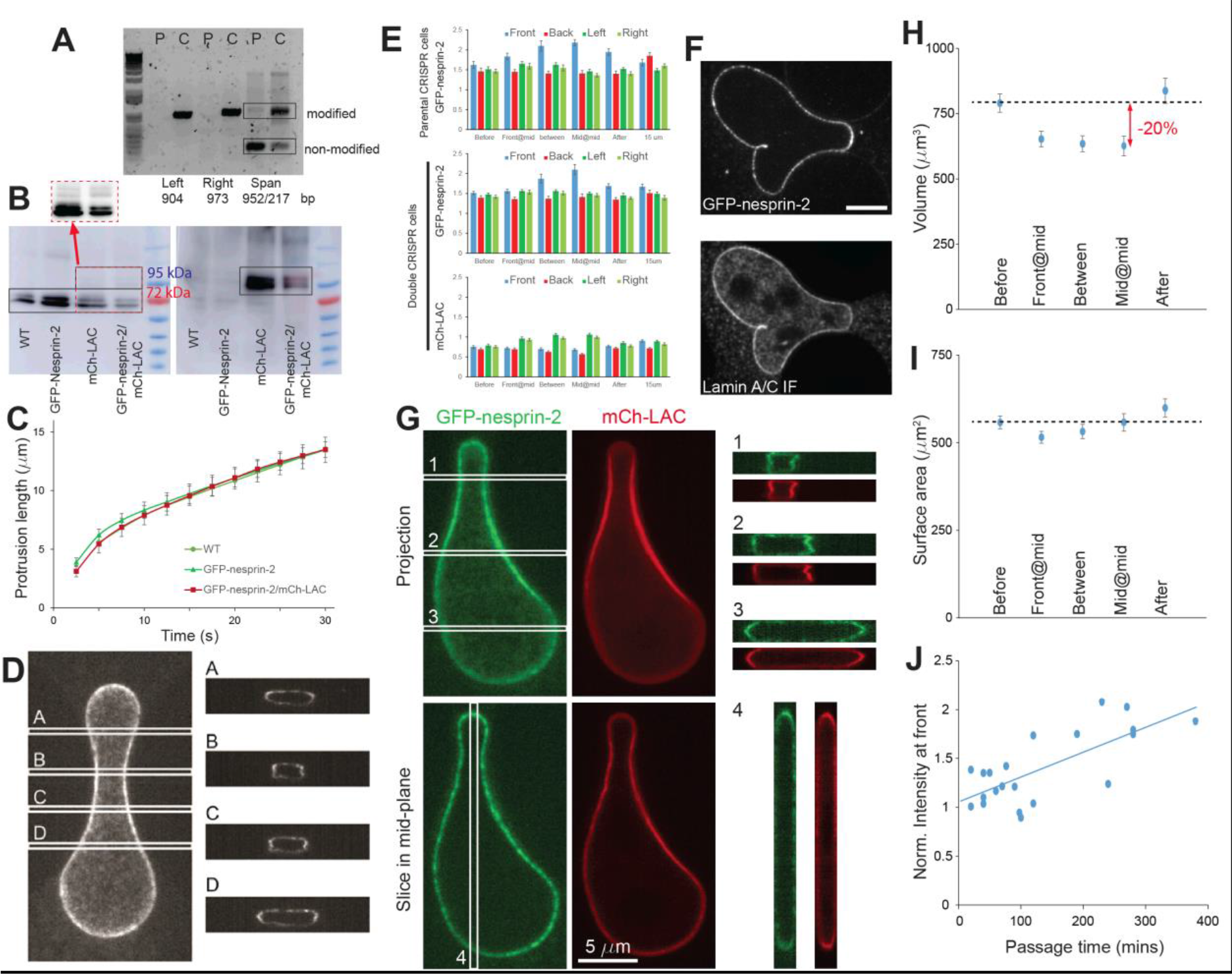
**A**, PCR to validate the insertion of mCh-Lmna. The gene sequences tested are shown in panel A. The sizes expected are labelled below the image. **B**, Western Blotting to validate the localization of bands at the expected size for Lamin A and C, and bands that correspond to the same protein with an additional 22 kDa, corresponding to the mCherry protein. The area outlined in red was reproduced at higher contrast to see the mCherry bands that are much less abundant than the wild-type lamins. **C**, Micropipette aspiration of the nucleus of cells bearing the CRISPR modification on nesprin 2 compared to cells bearing both this modification and the mCh insertion on lamin A/C. The additional mCh on lamins does not affect the rate of deformation in these devices. (Error bars correspond to the standard error of the mean. WT: 124 cells over 3 experiments; GFP-nesprin-2: 76 cells over 4 experiments, GFP-nesprin-2/mCh-LAC: 63 cells over 3 experiments.) **D**, Confocal imaging of a GFP-nesprin-2 cell migrating through a narrow constriction, along with reconstructions of 4 different cross-sections (right). **E**, Intensity measurements on the GFP-nesprin-2 cell line and the mCh-LAC/GFP-nesprin-2 cell line. The data is the same as the data presented in Figure 1 D and 2, but this data is not normalized. (Error bars correspond to the standard error of the mean. Parental: n=17 cells over 4 different experiments, double CRISPR: n=19 cells over 5 different experiments. In the 15 μm constrictions, parental: n=20 cells over 3 experiments, double CRISPR: n=27 over 6 experiments.) **F**, Immunofluorescence image of GFP-nesprin-2 cells migrating through a 2 μm constriction. **G**, Confocal imaging of a mCh-LAC/GFP-nesprin-2 cell migrating through a narrow constriction. Projections of the two fluorescent channels are shown (top), along with a single slice in the middle of the stack (bottom) and reconstructions of 4 different cross-sections (right). **H**, Volume of the nucleus as it is deformed through a constriction. The volume was approximated based on the cross-sectional area of nuclei expressing GFP-nesprin-2 and mCh-LAC imaged on an epifluorescence microscope, knowing that the channels are 5 μm tall and the nucleus fills the available space (see side projections in panel C). **I**, Surface area of the nucleus as it is deformed through a 2 μm constriction. The surface area is approximated from the cross-sectional area and perimeter of nuclei deformed through constrictions. **J**, Relationship between the nesprin accumulation and the passage time. The intensity at the front of the nucleus was normalized to the sides and plotted as a function of passage time through the constrictions (time between “Front@mid” and “Mid@mid”).

**Supplemental Figure 3:**
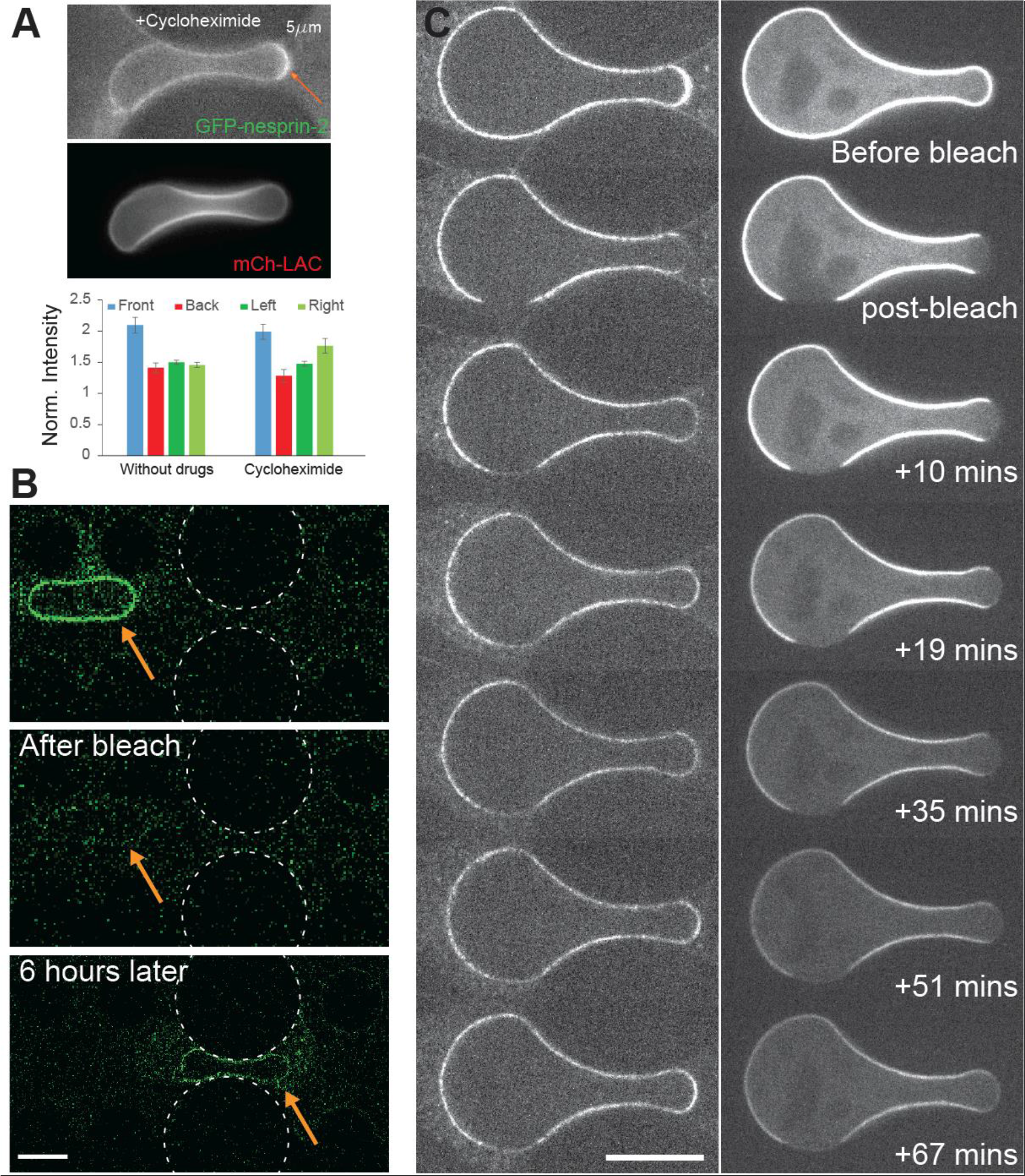
A) Quantification of the accumulation at the Mid@mid time point with cyclohexamide treatment (10 μg/ml). (Error bars correspond to the standard deviation, n=19 cells over 4 experiments in the “without drugs” conditions and n=6 cells over 2 experiments for the cycloheximide experiments. Comparison between front and back: without drugs p=3.27 × 10^−5^, Cycloheximide p=0.00125). **B**, Recovery of GFP-nesprin-2 signal in bleached nuclei. Images taken immediately before and after bleaching and then 6 hours later. After 6 hours the signal has only partially recovered: the turnover of GFP-nesprin-2 is longer than 6 hours. The signal in the bottom image has recovered uniformly around the periphery of a nucleus deformed in a constriction, indicating that newly synthesized protein is not selectively inserted at the front of the nucleus. **C**, Fluorescence recovery of bleached areas of GFP-nesprin-2 and mCh-LAC. Areas of GFP and mCh signal were bleached in cells migrating through 2 μm constrictions. Note that the GFP signal recovers to the extent that the signal is stronger at the front than the rest of the nuclear envelope, but the signal at the back has not recovered and neither has the lamin signal. (Error bars 10 μm.)

**Supplemental Figure 4:**
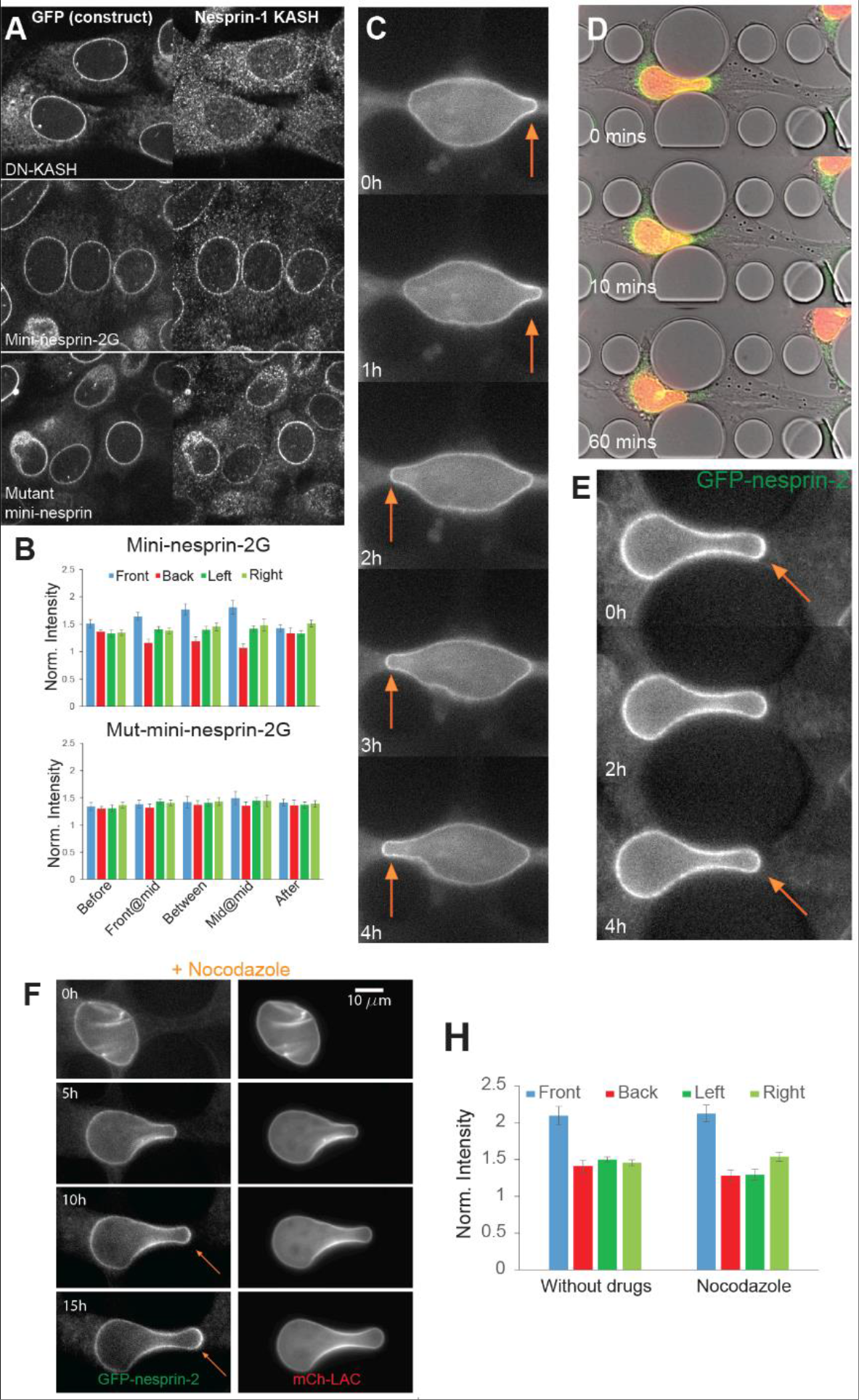
Actin is responsible for GFP-nesprin-2 accumulation. **A**, Cells expressing GFP constructs (left) labelled with an antibody for the KASH domain of Nesprin-1 result in displacement when the DN-KASH construct is expressed, but not when the mini-nesprin-2G or mutant-mini-nesprin-2G constructs are expressed. **B**, Quantification of the intensity around the periphery around the nucleus of cells expressing the mini-nesprin-2G and mutant constructs. This data is also represented in a different format in Figure 3C. (Error bars correspond to the standard error of the mean. Mini-nesprin-2G: n=10 cells over 4 different experiments; Mutant: n=11 cells over 4 different experiments.) **C**, Example of a cell that accumulates mini-nesprin-2G in one direction, but then loses this accumulation as the direction of migration is reversed. **D**, Addition of Cytochalasin D results in nuclei falling backward, even when adhesion at the front of the cell is not lost. (Green: GFP-nesprin-2, Red: mCh-LAC). **E**, GFP-nesprin-2 cells migrating through constrictions treated with Cytochalasin D. Upon addition of the drug the migration of the cells is drastically reduced. These cells no longer translocate their nuclei through constrictions. In some instances nuclei that are engaged into the constriction back out and partially lose their nesprin accumulation (orange arrow). **F**, Nocodazole treatment does not abolish accumulation of GFP-nesprin-2 (orange arrows) when cells migrate through the constrictions. **G**, Quantification of the accumulation at the Mid@mid time point upon nocodazole treatment (1μM). (Error bars correspond to the standard deviation, n=19 cells over 4 experiments in the “without drugs” conditions and n=6 cells over 2 experiments for the nocodazole treatment. Comparison between front and back: without drugs p=3.27 × 10^−5^, nocodazole p=1.67×10^−4^)

**Supplemental Figure 5:**
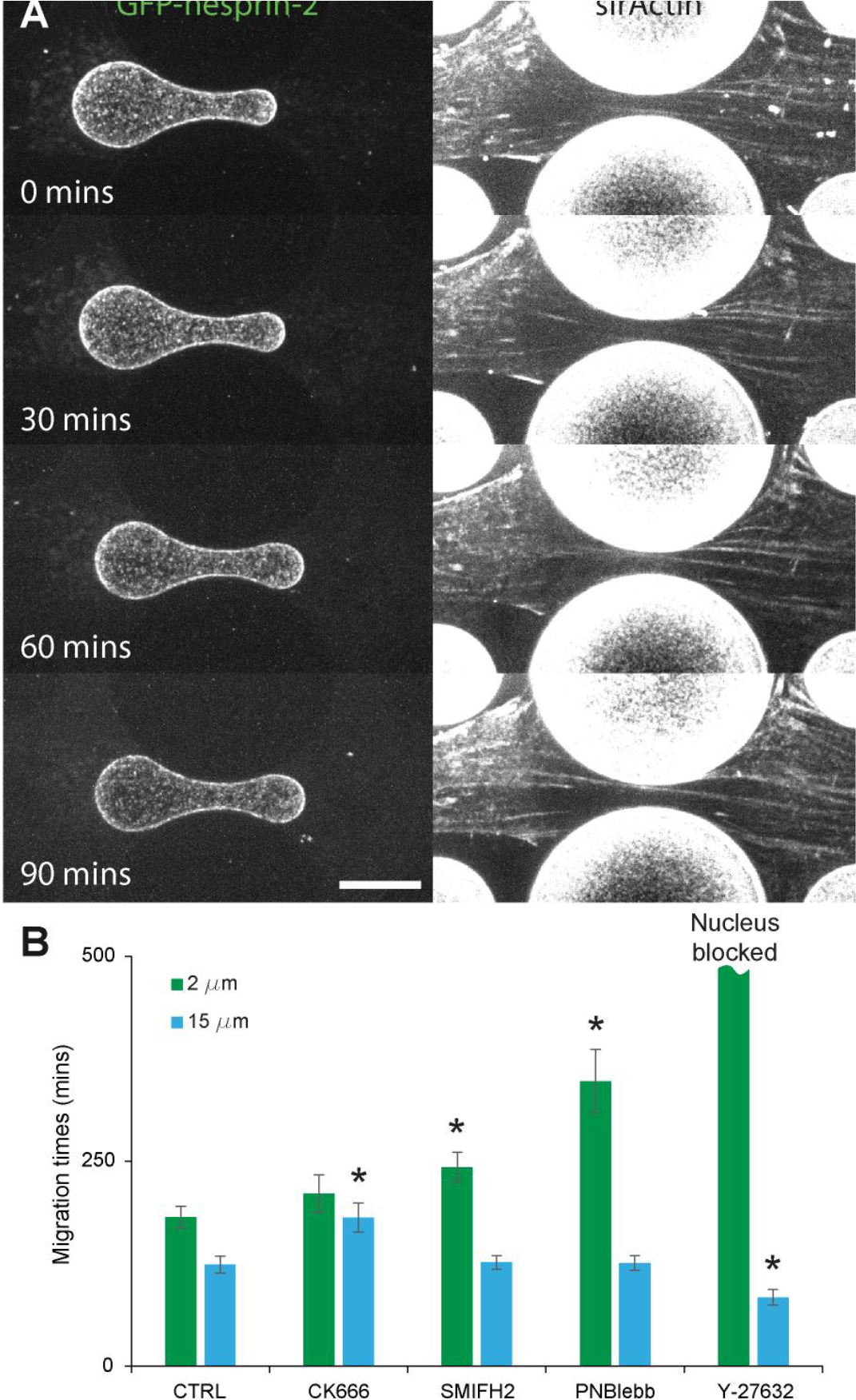
Actin organization during migration. **A**, Live cells were imaged on a spinning disk microscope during migration through a 2 μm constriction to obtain confocal images at different heights. The images at each timepoint were projected to obtain one image with minimal signal from the SiR-Actin absorbed onto the surfaces of the migration devices. **B**, Migration times of cells treated with actin or myosin-inhibiting drugs, or control (1:1000 DMSO). (All experiments performed over 3 different days; CTRL, 2 μm N=75, 15 μm N=46; CK666, 2 μm N=40, 15 μm N=50; SMIFH2, 2 μm N=54, 15 μm N=57; PNBlebb, 2 μm N=21, 15 μm N=47; Y-27632 experiments only quantified on day when a data point was obtained for the 2 μm constriction, 2 μm N=1, 15 μm N=25.)

## Supplementary Movies

**Supplementary movie 1:** Timelapse imaging of a GFP-nesprin-2 cell migrating through a 2 μm constriction. Top is the brightfield signal, bottom is the GFP signal. The data is the same as Figure 1C. Each frame is separated from the next by 10 minutes.

**Supplementary movie 2:** Timelapse imaging of a GFP-nesprin-2/mCh-LAC cell migrating through a 2 μm constriction. Top is the brightfield signal, middle is the GFP signal and bottom is the mCherry signal. The data is the same as Figure 2A. Each frame is separated from the next by 10 minutes. The time scale represents hours:minutes.

**Supplementary movie 3:** Timelapse imaging of a GFP-nesprin-2/mCh-LAC cell showing a FRAP experiment. Top is the brightfield signal, middle is the GFP signal and bottom is the mCherry signal. First frame is immediately before bleach and the second frame is immediately after bleach. The following images show recovery of fluorescence over five minutes.

**Supplementary movie 4:** Timelapse imaging of a cell expressing a GFP-mini-nesprin-2G construct migrating through a 3 μm constriction. Top is the brightfield signal, bottom is the GFP signal. The data is the same as Figure 4A. Each frame is separated from the next by 10 minutes. The time scale represents hours:minutes.

**Supplementary movie 5:** Timelapse imaging of a cell expressing a GFP-MUT-mini-nesprin-2G construct migrating through a 3 μm constriction. The GFP signal is shown. Each frame is separated from the next by 10 minutes. The time scale represents hours:minutes.

**Supplementary movie 6:** Timelapse imaging of a cell expressing a GFP-mini-nesprin-2G construct in microfluidic migration devices. The GFP signal is shown. The data is the same as Supplemental Figure 4C. Each frame is separated from the next by 10 minutes. The time scale represents hours:minutes.

**Supplementary movie 7:** Timelapse imaging of a GFP-nesprin-2 cell migrating through a 2 μm constriction. Left is the brightfield signal, right is the GFP signal. Each frame is separated from the next by 10 minutes. The time scale represents hours:minutes. Adhesion of the cell at the front in the brightfield image is demonstrated by a green circle and loss of adhesion is demonstrated by a red circle. Forward movement of the nucleus is indicated with a green arrow and when the nucleus falls back a red arrow is shown.

---

**Supplementary Table 1:**
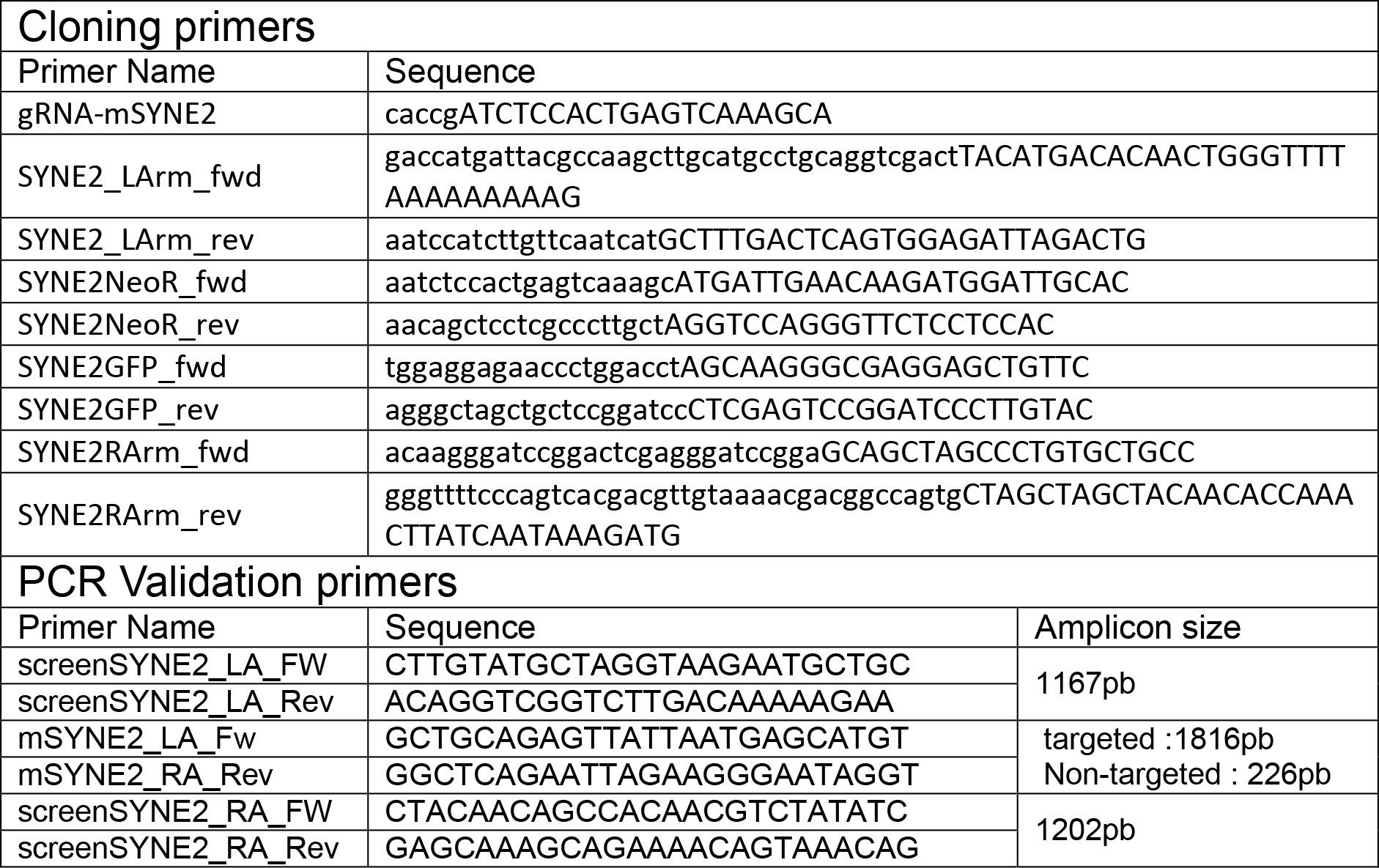
Primers for the Syne2-GFP modification.

**Supplementary Table 2:**
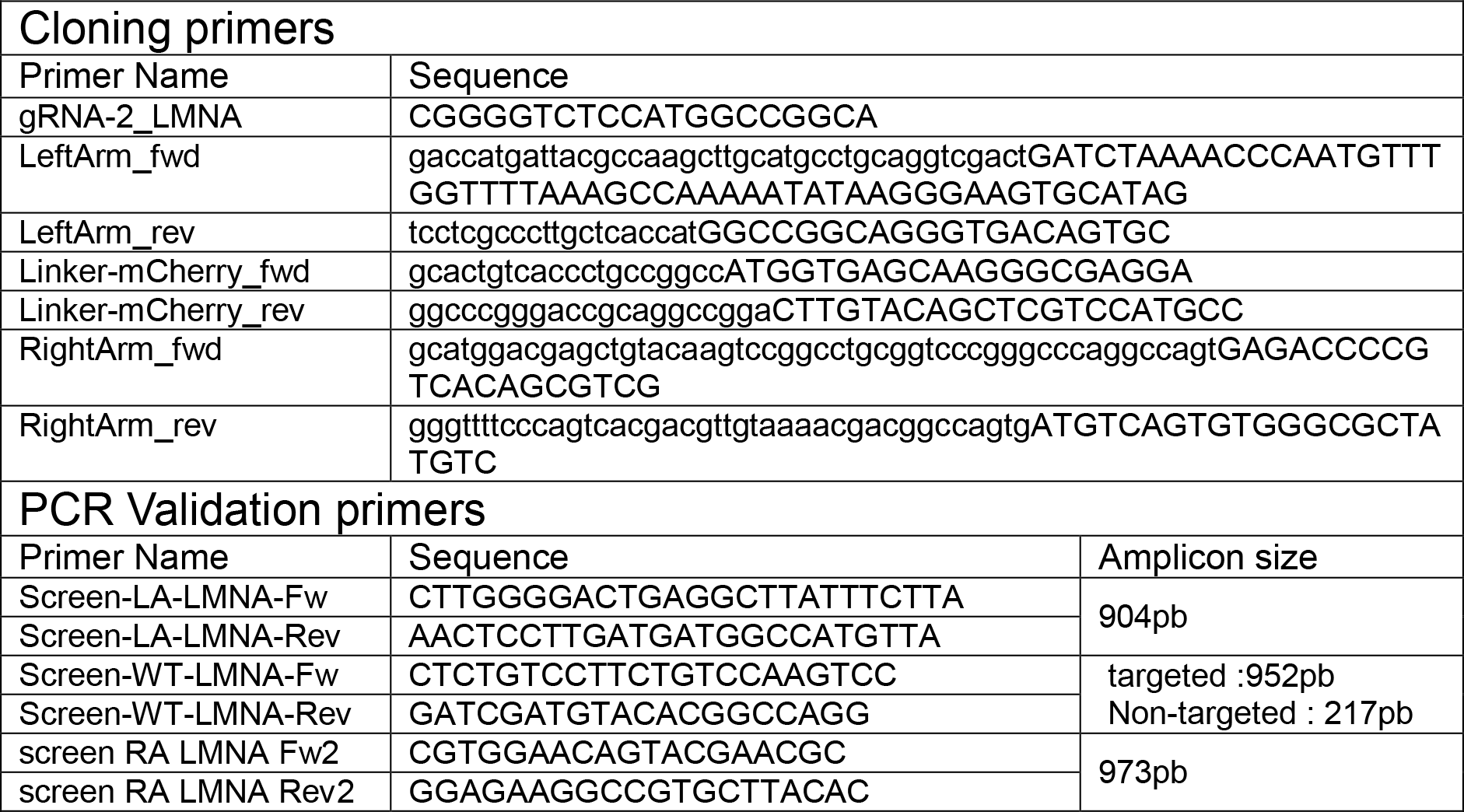
Primers for the Lmna-mCh modification.

**Supplementary Table 3:**
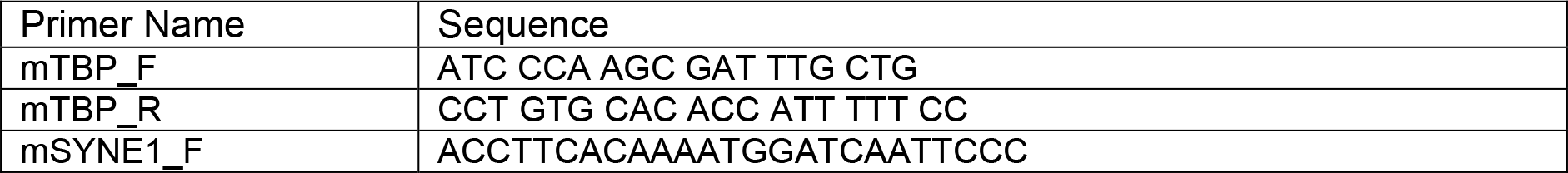
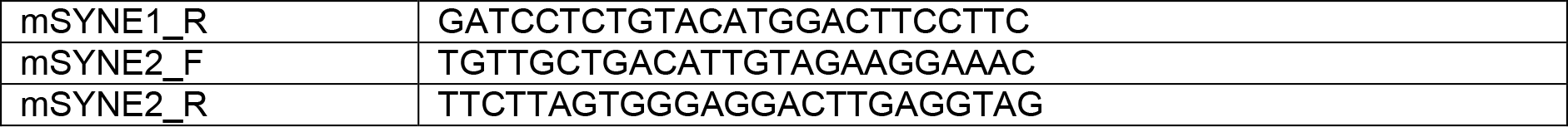
qPCR Primers.

**Supplementary Table 4:**
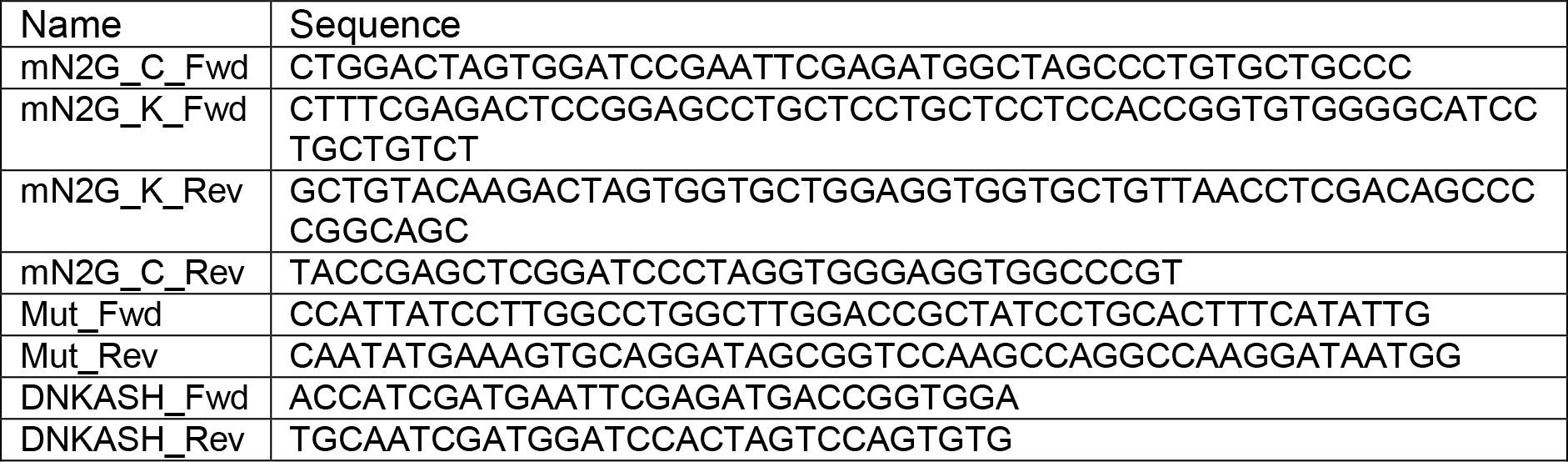
Cloning plasmids for NIH 3T3 cells.

